# Experienced meditators exhibit no differences to demographically-matched controls in theta phase synchronisation, P200, or P300 during an auditory oddball task

**DOI:** 10.1101/608547

**Authors:** JR Payne, O Baell, H Geddes, B Fitzgibbon, M Emonson, AT Hill, NT Van Dam, G Humble, PB Fitzgerald, NW Bailey

## Abstract

**Objectives:** Long-term meditation practice affects the brain’s ability to sustain attention. However, how this occurs is not well understood. Electroencephalography (EEG) studies have found that during dichotic oddball listening tasks, experienced meditators displayed altered attention-related neural markers including theta phase synchronisation (TPS) and event-related potentials (ERP; P200 and P300) to target tones while meditating compared to resting, and compared to non-meditators after intensive meditation interventions. Research is yet to establish whether the changes in the aforementioned neural markers are trait changes which may be observable in meditators irrespective of practice setting.

**Method:** The present study expanded on previous research by comparing EEG measures from a dichotic oddball task in a sample of community-based mindfulness meditators (n=22) to healthy controls with no meditation experience (n=22). To minimise state effects, neither group practiced meditation during / immediately prior to the EEG session.

**Results:** No group differences were observed in behavioural performance or either the global amplitude or distribution of theta phase synchronisation, P200 or P300. Bayes Factor analysis suggested evidence against group differences for the P200 and P300.

**Conclusions:** The results suggest that increased P200, P300 and TPS do not reflect trait-related changes in a community sample of mindfulness meditators. The present study used a larger sample size than previous research and power analayses suggested the study was suficiently powered to detect differences. These results add nuance to our understanding of which processes are affected by meditation and the amount of meditation required to generate differences in specific neural processes.

---

Mindfulness is a broad term that describes many practices, processes, and characteristics related to altering the capacities of attention, awareness, and acceptance (Van Dam et al., 2018). Mindfulness-based interventions (MBIs) may be an effective treatment for a range of psychological disorders (Keng, Smoski, & Robins, 2011), including anxiety (Miller, Fletcher, & Kabat-Zinn, 1995) and depression (Godfrin & van Heeringen, 2010; Ma & Teasdale, 2004), though the exact mechanisms of action are unclear (Goyal, Singh, Sibinga, & et al., 2014). There is additional evidence that mindfulness meditation, which involves deliberately focusing one’s attention on an aspect of present-moment experience in a nonjudgmental manner, can improve the psychological wellbeing of healthy people (Chambers, Gullone, & Allen, 2009; Gu, Strauss, Bond, & Cavanagh, 2015). Despite the potential importance of mindfulness in clinical and healthy populations, our understanding of the neurophysiological mechanisms underlying the benefits of mindfulness meditation and MBIs are restricted to theorised models which lack firm agreement (Schoenberg & Vago, 2019; Tang, Hölzel, & Posner, 2015). Mechanistic-based approaches allow for more precisely targeted interventions (Britton et al., 2018), so increased knowledge of these underlying mechanisms is valuable.

The most commonly implicated cognitive mechanism in contemporary mindfulness meditation known to be enhanced is attention (Gu et al., 2015; Tang et al., 2015). Indeed, developing the ability to control one’s attention is a common first step in traditional teachings of mindfulness meditation (Hölzel et al., 2011; Lutz, Slagter, Dunne, & Davidson, 2008), and studies have shown that experienced mindfulness meditators outperform non-meditators on attention-related cognitive tasks (Badart, McDowall, & Prime, 2018; Bailey et al., 2018; Cardeña, Sjöstedt, & Marcusson-Clavertz, 2015; Hodgins & Adair, 2010). Using brain imaging methods, such as functional Magnetic Resonance Imaging (fMRI) and electroencephalography (EEG), studies have found that experienced mindfulness meditators display enhanced activation in attention-related brain regions, such as the anterior cingulate and the prefrontal cortex when compared to controls (Allen et al., 2012; Bailey et al., 2018; Grant, Courtemanche, Duerden, Duncan, & Rainville, 2010; Tang et al., 2015; Tomasino & Fabbro, 2016). Furthermore, experienced meditators also display altered amplitudes of event-related potentials (ERP) related to attention (such as the P200 and P300) compared to people without meditation experience (Atchley et al., 2016; Cardeña et al., 2015; Lutz et al., 2009). By identifying neural markers of attention common to experienced meditators, mechanistic-based research could enable optimisation of the parameters of MBIs. This would allow the interventions to maximally target these markers in populations where improvements in attention may increase wellbeing.

There are a variety of differences in brain activity that have been found in experienced meditators, including state changes, which are dependent on the act of meditation, and trait changes, which can be observed independently of the act of meditation (Cahn & Polich, 2006). Trait changes in meditators are likely the result of repeated engagement of specific brain states over time (Davidson, 2005). Due to the overlap between state and trait changes, it is difficult to distinguish the two types of changes. Nonetheless, identifying broad attention-related neurophysiological trait changes is valuable, as they are likely to reflect stable changes and therefore are useful targets for research. Previous research has mainly focused on neural activity in very experienced meditators, or after an intensive meditation retreat (Atchley et al., 2016; Cahn & Polich, 2009; Cardeña et al., 2015; Delgado-Pastor, Perakakis, Subramanya, Telles, & Vila, 2013; Lutz et al., 2009). However, not all meditators engage in sustained, isolated practice, it is beneficial to observe a sample of meditators who live in the community but practice regularly. Exploring such a community sample rather than a committed, monastic-like sample, permits generalisable and translatable findings (these meditators will be referred to as “a community sample of meditators” henceforth).

A useful way of measuring brain changes in experienced meditators is EEG, as it is a non-invasive and relatively inexpensive technique which offers high temporal resolution. However, traditional ERP and phase locking EEG analysis methods to observe attention-related neural activity in meditators typically average neural activity measures to restricted time windows or specific electrodes (Klimesch, Sauseng, Hanslmayr, Gruber, & Freunberger, 2007; Koenig & Melie-garcía, 2010; Light et al., 2010). As fMRI studies have shown meditators to have altered distributions of neural activity including more neural activity in frontal areas such as the anterior cingulate and prefrontal cortex (Allen et al., 2012; Grant et al., 2010; Hölzel et al., 2011; Tang et al., 2015; Tomasino & Fabbro, 2016), EEG analysis methods which can detect not just amplitude changes at single electrodes but discriminate overall changes in neural response strength from alterations in the distribution of brain activity are warranted. The current study extends previous research which has focused on single electrode analyses to use a state of the art analysis technique that allow for separate comparisons of the amplitude and the distribution of neural activity (Koenig, Kottlow, Stein, & Melie-García, 2011). These have only been implemented in one study of meditation thus far (Bailey et al., 2018).

The ERP P300 and the less commonly studied P200 are two attention-related markers that have been observed with EEG in experienced meditators (Atchley et al., 2016; Lakey, Berry, & Sellers, 2011; Lutz et al., 2009; Slagter, Lutz, Greischar, Nieuwenhuis, & Davidson, 2009). The P300 is often observed in oddball paradigms when participants respond to a target stimulus that occurs infrequently among a series of frequently occurring standard stimuli (Arns, Jongsma, & Kessels, 2014). Increased P300 amplitudes to target tones during passive dichotic oddball tasks have been observed in experienced meditators while meditating compared to when not meditating (Delgado-Pastor et al., 2013) and studies have shown meditators to have increased P300 to target tones when asked to meditate throughout the task compared to non-meditators (Atchley et al., 2016; Jo, Schmidt, Inacker, Markowiak, & Hinterberger, 2016; Lakey et al., 2011; Sarang & Telles, 2006). Other studies have shown decreased P300 in meditators to distractor stimuli while meditating (Cahn & Polich, 2009) and during states of mind wandering (Barron, Riby, Greer, & Smallwood, 2011). Atchley et al. (2016) found that an increased P300 during dichotic listening tasks is a potential marker of long-term meditation experience in mindfulness meditators. In addition to the research showing differences in the P300, a study by Lutz et al. (2009) found that after 3-month retreat meditators, but not controls, displayed an increase in the P200 when attending to target tones during a dichotic listening task. The P200 is thought to be moderated by attention and associated with the start of an executive process when identifying unusual stimuli (Lindholm & Koriath, 1985). These studies suggest that P300 and possibly the P200 may be markers of neural attentional resource allocation that are altered and possibly increased to meet the demands of attention over time through meditation practice

A number of researchers have also suggested that alteration to theta (4–8 Hz) phase synchronisation (TPS) is an expected effect to be observed with extensive meditation training (Baird, Smallwood, Lutz, & Schooler, 2014; Cahn, Delorme, & Polich, 2013; Lutz et al., 2009; Slagter et al., 2009). In the context of a dichotic listening oddball task, TPS is the consistency of the angle of theta oscillations time-locked to the target stimulus onset across trials (a measure which is independent of the theta oscillation’s amplitude) (Klimesch, Doppelmayr, Schimke, & Ripper, 1997). Increased TPS is associated with enhanced attention and meditation across different types of practice (Aftanas & Golocheikine, 2001; Baijal & Srinivasan, 2010; Cahn et al., 2013; Lutz et al., 2009). TPS has been found to be increased when responding to target tones during a dichotic listening task in meditators whilst meditating compared to when resting (Cahn et al., 2013). Lutz et al. (2009) found that meditators showed a stronger negative correlation between TPS and response variability than non-meditators who did not attend the retreat. Higher TPS values were associated with lower variability of responses, a measure which suggests increased stability of attention, and which the authors interpreted as increased sustained attention. However, in Lutz et al. (2009)’s study, participants were instructed to complete the task as a focused attention meditation practice after briefly meditating and attending an intensive three-month meditation retreat. These studies suggest that TPS may be involved with the state of meditation and could be a trait marker of extensive meditation practice over time. Research has not explicitly determined whether alterations in TPS, the P300 and P200 are a trait effect among experienced meditators. Additionally, previous research has often observed experienced meditators who live a monastic lifestyle who engage in regular intensive retreats, rather than experienced meditators who live in the community and practice regularly (Thomas & Cohen, 2014).

The aim of the current study was to assess whether a community sample of experienced meditators would show trait alterations in attention-related neurophysiological markers such as P200, P300, and TPS compared to demographically-matched control participants with no meditation experience. To achieve this aim participants were asked to complete an active dichotic listening oddball task (replicating the task used by Lutz et al. (2009)), which is a putative measure of neural activity related to sustained attention. Overall, it was hypothesised that meditators would show enhancements to neural markers of sustained attention. Specifically, it was hypothesised that meditators would show 1) increased neural strength in the P200, P300, and TPS to attended target tones compared to non-meditators. It was also hypothesised 2) that meditators would display more frontal distribution of P200, P300 and TPS than non-meditators. Furthermore, 3) it was hypothesised that there would be a negative correlation between increased TPS and reduced response variability to attended target tones. Lastly, 4) it was hypothesised that meditators would respond more quickly and accurately and show less variability in reaction time than non-meditators.

## Method

### Participants and Self-Report Data

A sample of 26 meditators and 27 healthy control non-meditators were recruited after responding via phone call or email to community advertising at universities, hospitals, meditation organisations, and on social media. To be considered as an experienced meditator, participants were required to have had at least two years of meditation experience and have been practising for a minimum of two hours per week over the last 3 months (the sample had a mean of 9.09 years of meditation experience and 6.35 hours of current practice per week). Meditation was defined by Kabat-Zinn’s definition: “paying attention in a particular way: on purpose, in the present moment, and nonjudgmentally” (Kabat-Zinn, 1994). This definition included participants who practice both focused attention (FA), which involves deliberate attention on a specific object, such as the breath and open monitoring (OM) meditation, which involves simple awareness without a specific focus besides awareness itself (Cahn & Polich, 2006; Lutz et al., 2008). Trained meditation researchers (JRP, OB, HG, NWB) screened and interviewed participants to ensure the participants practice fit the criteria. Healthy control participants were deemed eligible if they had less than two hours of lifetime meditation experience. Screening uncertainties were resolved by consensus between two researchers including the principal researcher (NWB).

Both meditator and control participants were considered ineligible to participate if they were currently taking psychoactive medication; had experienced brain injury; had previously been diagnosed with a psychiatric or neurological condition; or met the criteria for any drug, alcohol or neuropsychiatric disorders as measured by the Mini International Neuropsychiatric Interview (MINI) DSM-IV (Hergueta, Baker, & Dunbar, 1998). Participants who scored above the moderate range for anxiety or depression in the Beck Anxiety Inventory (BAI) (Steer & Beck, 1997) or Beck Depression Inventory II (BDI-II) (Beck, Steer, & Brown, 1996) were also excluded. All participants were between 20 and 60 years of age, see Table 1.

**Table 1.** Participant Demographic and Behavioural Data

|  | Meditators | Controls | Statistics |
| --- | --- | --- | --- |
|  | *M (SD)* | *M (SD)* |  |
|  | (n = 22) | (n = 22) |  |
| Age | 33.27 (13.13) | 31.82 (11.12) | t(42) = 1.162, p = 0.694 |
| Gender (M/F) | 8/14 | 7/15 | Chi-square = 0.10, p = 0.750 |
| Years of Education | 15.41 (3.30) | 17.34 (2.64) | t(42) = 2.144, p = 0.038 |
| Meditation Experience (years) | 9.09 (7.39) | 0 |  |
| Current Time Meditating Per Week (hours) | 6.35 (3.26) | 0 |  |
| BAI score | 6.45 (6.5) | 5.23 (4.51) | t(42) = -0.727, p = 0.471 |
| BDI score | 3.36 (4.41) | 4.41 (5.10) | t(42) = 0.727, p = 0.471 |
| FFMQ score | 154.77 (12.54) | 132.59 (14.71) | t(42) = -5.380, p < 0.001** |

*Note:* n = total number of participants in each group; \*\* = p < 0.01; SD = standard deviation; BDI = Beck Depression Inventory; BAI = Beck Anxiety Inventory; FFMQ = Five Facet Mindfulness Questionnaire.

Before participants underwent EEG recording, participants provided their gender, age, years of education, handedness, and meditation experience (total years of practice, frequency of practice, and usual length of meditation session). Participants also completed the Five Facet Mindfulness Questionnaire (FFMQ) (Baer, Smith, Hopkins, Krietemeyer, & Toney, 2006), BAI, and BDI-II. Table 2 summarizes these measures. All participants provided written informed consent prior to participation in the study. Ethical approval of the study was provided by the Ethics Committees of Monash University and the Alfred Hospital.

**Table 2.**
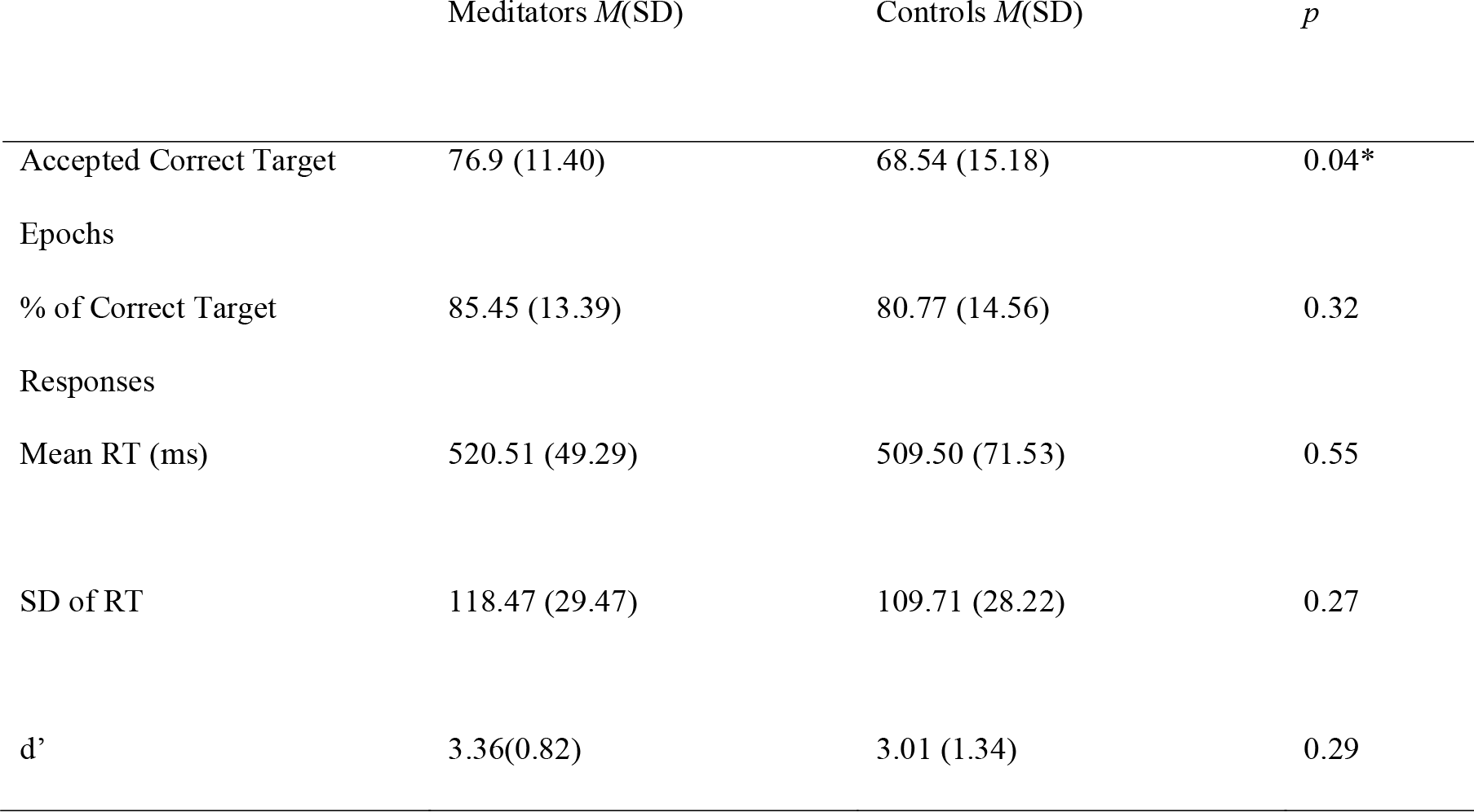
Participant Behavioural Data

One control was excluded from the study after scoring above the moderate anxiety range on the BAI. Two controls were excluded due to technical faults while recording. Additionally, two controls and four meditators were excluded from analysis after providing less than 38 accepted target stimuli EEG epochs. The final sample included 22 meditators aged between 20 and 58 years (14 men, 8 women, Mage = 33.27 years, SDage = 13.13 years), and 22 healthy controls aged between 23 and 60 (15 men, 7 women, Mage = 31.82 years, SDage = 11.12 years).

### Procedure and Task

Participants performed a Go/Nogo task (data not yet published) followed by the dichotic listening oddball task. The dichotic listening oddball task was a replication of the “focused attention” version used by Lutz et al. (2009). The task involved four blocks where the participants heard tones in both ears and were instructed to focus on the tones in one ear only (i.e. the attended ear) and ignore the tones from the other ear (i.e. the not attended ear). Each block began with instructions as to which ear to attend to (left or right). Participants discriminated between two kinds of tones in the attended ear, frequent standard tones and rare target tones. Target tones were at a lower or higher hertz (Hz) than the standard tone, and participants were instructed to respond as quickly and accurately as possible to target tones. The instructions included examples of standard and target tones and instructed participants to keep their eyes fixated on a cross on the computer screen to minimise head movements during the task. All blocks began with 75 auditory practice stimuli followed by 350 real stimuli. Each stimulus was 80 decibels, with a duration of 60 milliseconds (ms) and a rise and fall time of 10 ms. The interstimulus interval (ISI) was randomised between 700 and 1100 ms (e.g. 767ms, 1021ms, 894ms). Each block contained 300 standard stimuli (one ear, 1000Hz; *n* = 150; the other ear, 500Hz; *n* = 150) and 50 target stimuli (one ear, 1050Hz; *n* = 25; the other, 475Hz; *n* = 25) (Lutz et al., 2009). The high tones (1000Hz standard; 1050Hz target) and low tones (500Hz standard; 475Hz target) were presented randomly in each ear across blocks, and blocks were presented in a random order across participants. The total task time was approximately 30 minutes. The participants then completed an attentional blink paradigm and were administered transcranial magnetic stimulation concurrent with EEG (data not yet published, note that the participants in this study comprise a separate sample to those in Bailey et al. 2018 and 2019).

### Electrophysiological Recording and Pre-Processing

64-channel EEG data were continuously recorded throughout the tasks using an EEG Ag/AgCI Quick-Cap amplifier (Compumedics, Melbourne, Australia). Data were integrated into Neuroscan Aquire software and a SynAmps 2 amplifier (Compumedics, Melbourne, Australia). Data recordings were sampled at 1000Hz and online bandpass filtered from 0.05 to 200Hz (24dB/octave roll off). Each electrode was referenced to a reference electrode positioned between CPz and Cz. The impedances of the electrodes were maintained at less than 5kΩ. Eye movements were recorded from one supraorbital electrooculography electrode above the left eye in line with the pupil. EEG were analysed offline in MATLAB version number R2017b (The Mathworks, Inc.). Data were pre-processed in EEGLAB (Delorme & Makeig, 2004) and FieldTrip (Oostenveld, Fries, Maris, & Schoffelen, 2011) for frequency analysis to compute inter-trial coherence in the theta frequency (4-8Hz) (TPS). Second order Butterworth filtering was applied to the data with a bandpass filter from 1-80Hz and a bandstop filter 47-53Hz to reduce line noise. Neural activity was then epoched to the onset of the task stimuli (−100 to 1000ms). Artefacts and bad electrodes were removed from analysis based on an automated procedure applied in EEGLAB and eye movement and muscle activity were removed by independent component analysis (data processing steps described in supplementary materials) (Delorme & Makeig, 2004).

Epochs from target trials used in the behavioural analysis (including both attended and not-attended target trials) from each condition and participant were averaged separately for ERP analyses. TPS was quantified through the calculation of a phase-locking factor (PLF) value (Lachaux, Rodriguez, Martinerie, & Varela, 1999; Ueno et al., 2009). PLF values range from 0 to 1, whereby value 1 represents perfectly correlated phase differences between trials, and 0 represents completely uncorrelated phase differences (Ueno et al., 2009; Varela et al., 2001). TPS computations are described in the supplementary materials.

## Statistical Comparisons

### Behavioural and Demographic Comparisons

Between-group comparisons of the behavioural and demographic data were performed using SPSS 23. Independent samples t-tests compared age, BAI, BDI, FFMQ, years of education, number of total accepted attended target epochs, percentage of correct responses, standard deviation (SD) of reaction time (RT) to target tones, mean RT and sensitivity index (d’) between groups.

### EEG Comparisons

EEG data comparisons of both ERPs and TPS between meditators and non-meditators were performed using the Randomised Graphical User Interface (RAGU) method (Koenig et al., 2011). RAGU compares scalp field differences over all epoch time points and electrodes using rank order randomisation statistics with no preliminary assumptions about time windows and electrodes to analyse (Koenig et al., 2011). Prior to conducting primary tests, a Topographical Consistency Test (TCT) was conducted to confirm consistent distribution of scalp activity within each group and condition. A significant TCT test suggests that potential between-group differences in the Global Field Power (GFP) test are due to real group differences, instead of variation within one of the groups (Koenig & Melie-garcía, 2010). RAGU allows for comparisons of global neural strength (independent of the distribution of activity) with the GFP test. The GFP is an index of the total voltage differences across all channels; it is equivalent to the standard deviation across all channels at every time point (Habermann, Weusmann, Stein, & Koenig, 2018). The GFP compares differences between groups from the real data against the randomised permutation data to determine specific time periods where groups significantly differed in neural response strength. RAGU also allows for comparisons of distribution of neural activity with the Topographic Analysis of Variance (TANOVA) (normalised for the amplitude of neural activity, and thus distribution comparisons are independent of differences in global amplitude) and Topographical Analysis of Covariance (TANCOVA) which performs the same operations as TANOVA except it compares neural data to a linear predictor instead of between-group comparisons.

TPS values were compared with Root Mean Square (RMS) and TANOVA tests (to separately compare overall neural response strength and distribution of neural activity, respectively). It should be noted that when phase synchronisation comparisons are computed with RAGU, the average reference is not computed on the transformed data (the average reference was computed prior to the transforms). As such, the test is a comparison of the RMS between groups, a measure which is a valid indicator of neural response strength in the phase synchronisation domain. In other respects, the statistic used to compare RMS between groups is identical to the GFP test described in the previous paragraph.

RAGU controls for multiple comparisons in space and time through global duration statistics which calculate the periods of significant effects within the epoch that are longer than 95% of significant effects in the randomised data with the alpha level at 0.05 (Koenig et al., 2011). The recommended 5000 randomisation permutations were conducted with an alpha of *p* = 0.05. Also, global count statistics and area under the curve statistics of significant time points were observed to confirm adequate control for multiple comparisons in the time dimension. For more in-depth information about RAGU and its analyses, please refer to Koenig et al. (2011), Koenig and Melie-garcía (2010) and Habermann et al. (2018).

### Comparisons of Event-Related Potentials

GFP and TANOVA tests were used to conduct repeated measures Analysis of Variance (ANOVA) design statistics, examining 2 group (meditators vs controls) x 2 condition (attended target vs not attended target) comparisons for ERP data from −100 to 1000 ms surrounding stimulus onset of the target tone. GFP and TANOVA tests were averaged between 150 to 300 ms (P200 period) (Lindholm & Koriath, 1985) and 300 to 500 ms (P300 period) (Polich, 2007) after the attended target tone to make direct comparisons with Lutz et al. (2009).

### Comparisons of Theta Phase Synchronisation

To compare TPS between the groups RMS and TANOVA tests were used to conduct repeated measures ANOVA design, examining 2 group (meditators vs controls) x 2 condition (attended target vs not attended target) comparisons for TPS data from 0 to 900 ms surrounding the target tone. RMS and TANOVA tests were averaged from 150 to 300 ms and from 300 to 500 ms after the attended target tone to make direct comparisons with Lutz et al. (2009).

### Theta Phase Synchronisation Correlations with SD of RT

Covariance GFP tests were conducted using participants SD of RT as a covariate to measure the relationship between response variability and TPS (0 to 900ms after attended tones). Covariance GFP tests were averaged between 300 to 500 ms after the attended target tone to make direct comparisons with Lutz et al. (2009).

### Single Electrode Replication Comparisons

In addition to the RAGU analysis, traditional single electrode comparisons were conducted for comparison with previous research. ERP data from electrodes Fz, FCz, and Cz had activity averaged during the P200 window (150-300ms) and from electrodes Fz, FCz, Cz, CPz, and Pz, during the P300 window (300-500ms). TPS data from electrode Fz was averaged between 150-300ms and 300-500ms. The averaged windows were calculated for both attended and not attended responses. The statistical program JASP (JASP Team, 2019) was used to perform single electrode analyses. Bayesian as well as frequentist repeated measures ANOVAs were used to conduct a 2 group (meditators vs controls) x response (attended vs not attended) x 3 electrode (Fz, FCz, Cz for the P200) and 5 electrode (Fz, FCz, Cz, CPz, Pz, for the P300) comparison for P200 and P300 respectively, in direct replication of Lutz et al. (2009) and Atchley, et al. (2016). Similarly, a Bayesian as well as frequentist repeated measures ANOVA was used to conduct a 2 group (meditators vs controls) x response (attended vs not attended) comparison for TPS at Fz during the 150-300ms window and the 300-500ms windows separately in direct replication of Lutz et al. (2009). Sphericity violations were resolved through the Greenhouse-Geisser correction (Greenhouse & Geisser, 1959). Furthermore, Bayesian and Pearson’s correlations were conducted between TPS activity at the Fz electrode during the 300-500ms window and participant’s SD of RT. Bayes Factor analyses were used to calculate the probability of the null hypothesis in contrast to the alternative hypothesis where null results were found (Rouder, Morey, Verhagen, Swagman, & Wagenmakers, 2017). The suggested comparison between models containing a hypothesized effect to equivalent models stripped of the effect (excluding higher order interactions) was performed for these analyses.

### Power Analysis

A power analysis was calculated to determine if any null results may be explained by a lack of power. Power analysis for TPS was conducted via extraction of t values from Lutz et al. (2009) (t = 5.8 with n = 13 provides Cohen’s d = 0.85) as the current study directly replicated their task. Although the current study was a cross-sectional rather than longitudinal in design, Lutz et al. (2009) did not report effect size or t-statistics for between-group comparisons at their second timepoint, and it was assumed meditators TPS before and after the intervention would provide the similar effects to the between group comparisons in the current study. Power analysis for P300 was extracted from the F-values of Atchley et al. (2016) (F = 4.56, n = 42, d = 0.659) when comparing meditators to non-meditators during a dichotic oddball listening task during a breath counting task vs during a tone counting task. Power analysis for P200 was extracted from the t-values of Lutz et al. (2009) (t = 2.8, n = 13, d = 0.64) when comparing meditators P200 before and after a 3-month retreat. The values were entered into Gpower (Faul, Erdfelder, & Buchner, 2007) to compute post-hoc power for TPS comparisons in an independent samples t-test with a one-way tail and alpha of 0.05 based on the present study’s sample size (n = 44). Similarly, to compute post-hoc power for P300 comparisons, a repeated measures ANOVA was computed with a one-way tail and alpha of 0.05 based on the present study’s sample size (n = 44). Furthermore, power was computed for a correlational design, using the Pearson’s R-value reported by Lutz et al. (2009) for the correlation between SD of RT and TPS (r = −0.40) and alpha of 0.05 based on the current study’s sample size (n = 44).

## Results

### Demographic and Behavioural Data

Meditators and controls did not significantly differ in age, BDI, or BAI (all p > 0.4). However, meditators reported significantly higher FFMQ scores than controls (controls: M = 132.59, SD = 14.71, meditators: M = 154.77, SD = 12.54), t(42) = −5.38, *p* < .001, 95% CI [−30.50, −13.86]. Controls reported significantly more years of education than meditators (controls: M = 17.34, SD = 2.64, meditators: M = 15.41, SD =3.30), t(42) = 2.144, p = 0.038. See Table 1.

Data met assumptions for the statistical tests used, see supplementary materials. The independent samples t-test revealed a significant main group effect of total accepted attended target epochs (t(42) = −2.06, *p* = 0.045, 95% CI [−16.53, −0.19]), no main effect of group in percentage of correct target tones, d’ (t(42) = −1.06, *p* = 0.29, 95% CI [−1.03, 0.32]), mean RT (t(42) = −0.59, *p* = 0.55, 95% CI [−48.39, 26.36]). and SD of RT to target tones (t(42) = 1.01, *p* = 0.32, 95% CI [−8.80, 26.31]). A summary of these results can be seen in Table 2.

### Neural Data

As RAGU uses randomisation statistics (permutation tests) with rank ordering to obtain statistical significance, normality and other assumptions are not required to be met (Koenig et al., 2011). TCT tests were conducted to test for within group consistency prior to GFP tests.

### No Difference in Event-Related Potentials

The TCT showed topographical consistency of neural activity for both groups during target attended and not attended trials across the majority of time periods between −100 and 1000ms surrounding the presentation of a target stimulus, including during most of the P200 period (150-300ms) and all of the P300 period (300-500ms) (See Figure S1 in supplementary materials). The 2×2 (Group x Condition) GFP test showed no significant main effect of group in ERP in a period −100 and 1000ms surrounding target presentation following attended and not attended target trials (global count *p* = 0.67, see Figure 1A). However, there was a significant main effect of condition in this period, with higher GFP in the attended condition (global count *p* < 0.01, see Figure 1B). There was no significant interaction effect in this period (global count *p* = 0.76, see Figure 1C). The GFP test averaged across the 150 and 300 ms (P200 period) showed no significant main effect of group in GFP (p = 0.29), there was a significant effect of condition GFP (p = 0.04), and no significant interaction effect of group (meditator versus control) and condition (attended versus not attended) (*p* = 0.39). See Figure 2A. The GFP test averaged across the 300 and 500 ms (P300 period) showed no significant man effect of group in GFP (p = 0.52), there was a significant effect of condition GFP (p < 0.01), and no significant interaction effect of group (meditator versus control) and condition (attended versus not attended) (*p* = 0.61). See Figure 2B.

**Fig 1.**
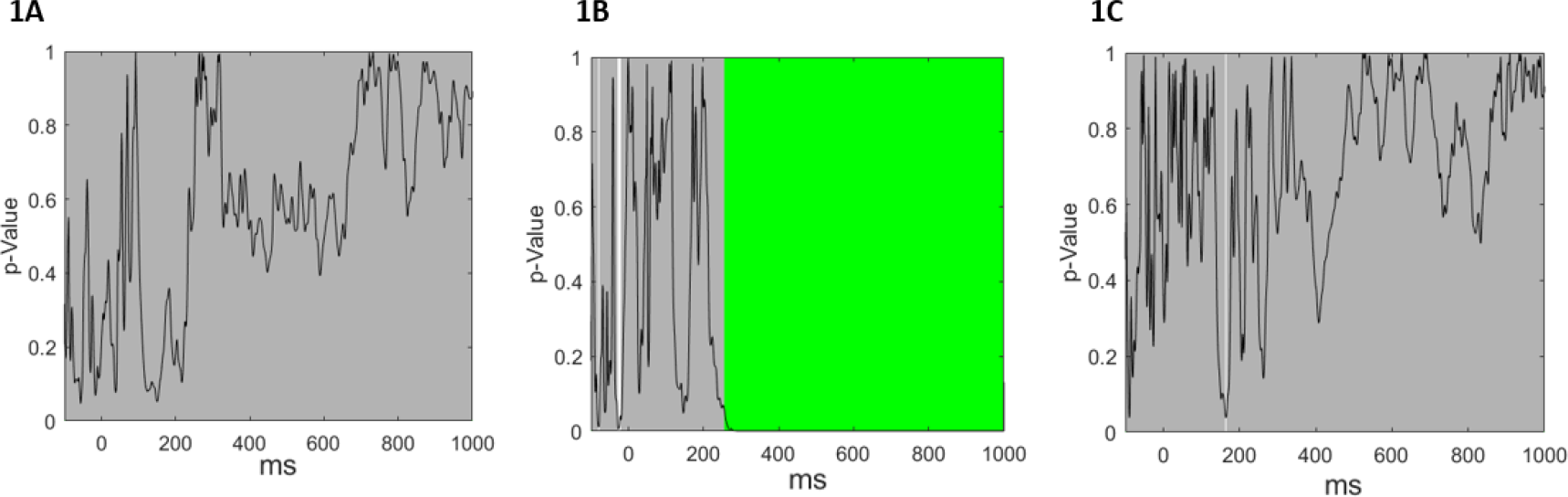
(**A**) No significant main effect of group was present in ERP GFP −100 and 100 surrounding stimulus presentation (global count *p* = 0.67). The black line shows *p-*value across the epoch. (**B**) There was a significant effect of condition in ERP across this window (global count *p* < 0.01). Areas marked in green indicate the duration of a significant effect that was greater 95% of significant effect in the randomised data, α = .05. Duration of significance within the epoch ranged from 253ms to 1000ms. (**C**) There was no significant interaction effect (group x condition) in this period (global count *p* = 0.76).

**Fig 2.**
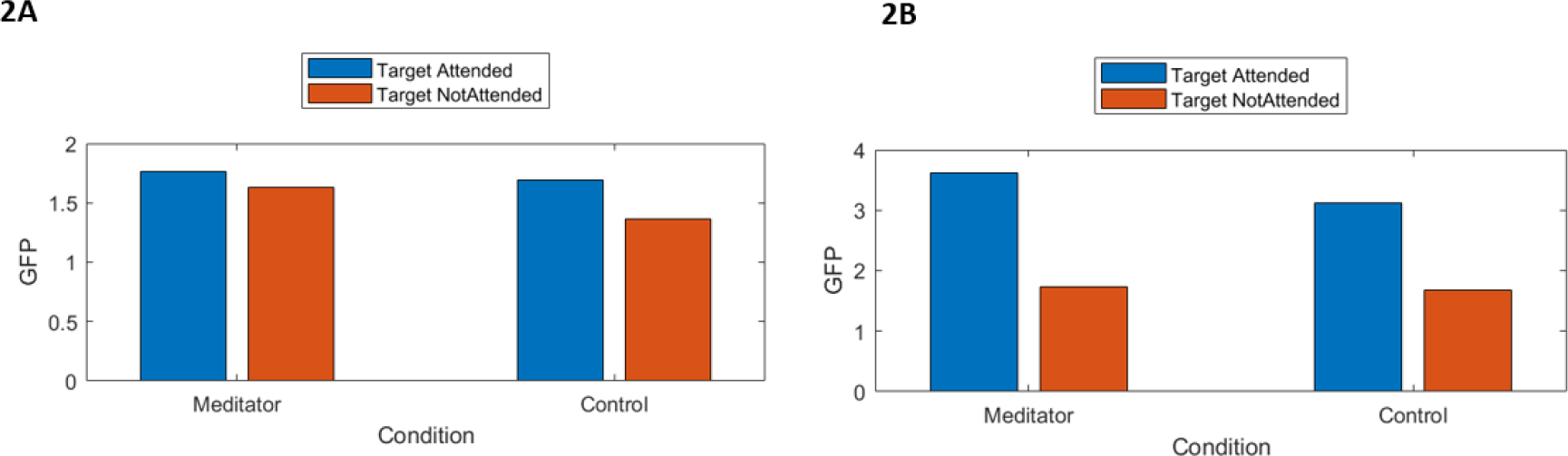
(**A**) There was no significant main effect of group in ERP GFP averaged across the P200 window (150-300ms) (p = 0.29). There was a significant effect of condition in ERP GFP averaged across the P200 window (p = 0.04). There was no significant interaction effect (group x condition) effect in ERP GFP averaged across the P200 window (p = 0.39). (**B**) There was no significant main effect of group in ERP GFP averaged across the P300 window (300-500ms) (p = 0.52). There was a significant effect of condition in ERP GFP averaged across the P300 window (p < 0.01). There was no significant interaction effect (group x condition) effect in ERP GFP during trials during the P300 window (p = 0.61).

The 2×2 (Group x Condition) TANOVA test showed no significant main effect of group ERP distribution in any periods −100 and 1000ms surrounding target presentation (global count *p* = 0.89, see Figure 3A). However, there was a significant main effect of condition from 199ms to 1000ms, with a distribution difference in the attended condition (global count *p* < 0.01, see Figure 3B). There was no significant interaction effect (group x condition) in this period (global count *p* = 0.76, see Figure 3C). The TANOVA test averaged across the 150 to 300 ms (P200 period) showed no significant main effect of group in distribution (p = 0.83), there was a significant effect of ERP distribution between conditions (p < 0.01), and no significant interaction effect of group (meditator versus control) by condition (attended versus not attended) (*p* = 0.42). See Figure 4A. The TANOVA test averaged across the 300 to 500 ms (P300 period) showed no significant main effect of group in ERP distribution (p = 0.70), there was a significant effect of ERP distribution between conditions (p < 0.01), and no significant interaction effect of group (meditator versus control) by condition (attended versus not attended) (*p* = 0.79). See Figure 4B.

**Fig 3.**
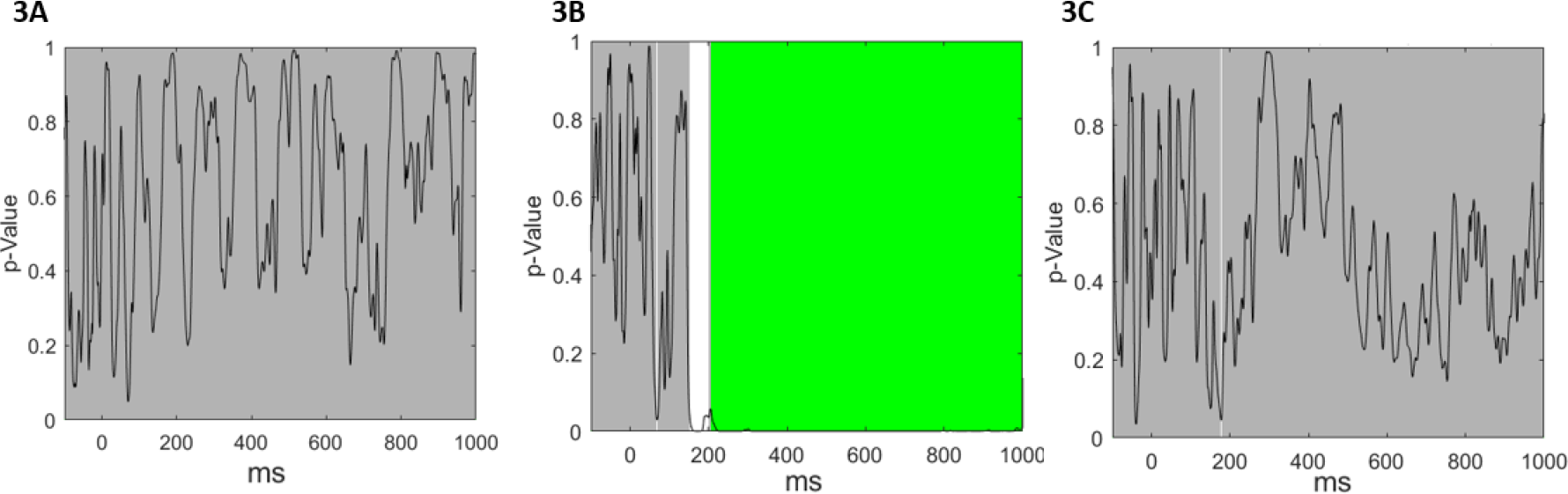
(**A**) The TANOVA revealed that there was no significant main effect of group in ERP distribution (global count *p* = 0.89). (**B**) There was a significant effect of condition in ERP distribution across this window (global count *p* < 0.01). Areas marked in green indicate the duration of a significant effect that was greater 95% of significant effect in the randomised data, α = .05. Duration of significance within the epoch ranged from 199ms to 1000ms. 1000ms (**C**) There was no significant interaction effect (group x condition) in this period (global count *p* = 0.76).

**Fig 4.**
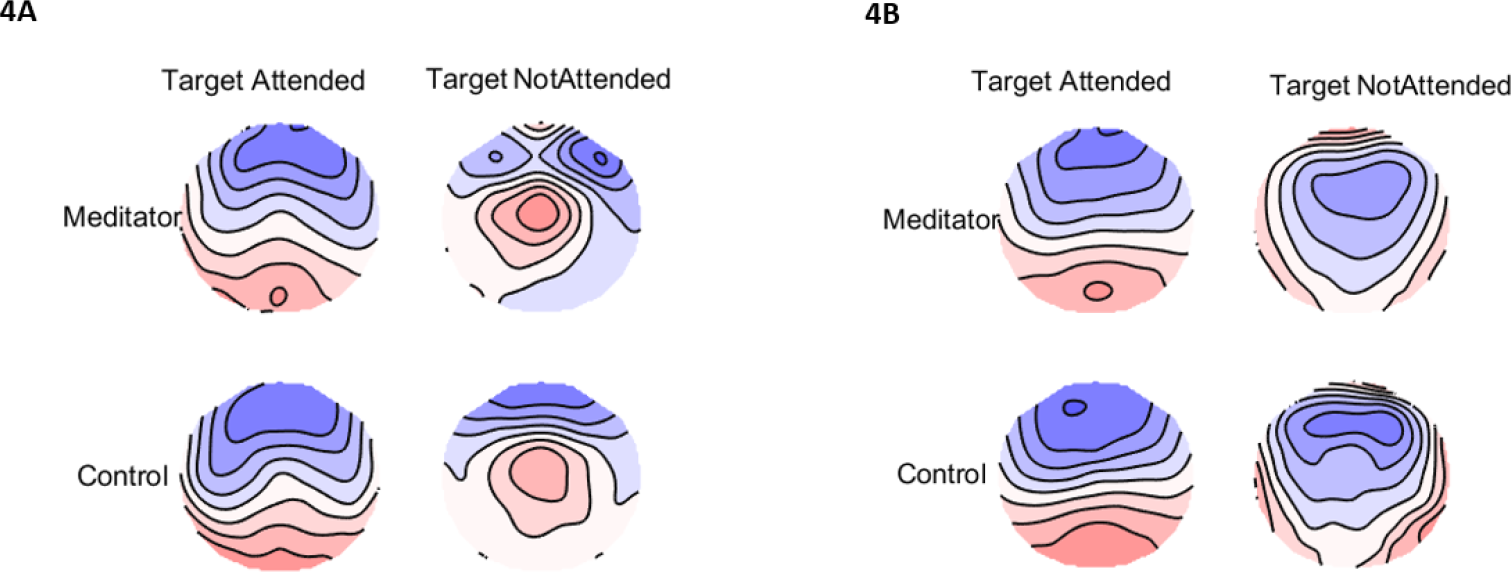
(**A**) There was no significant main effect of group in ERP distribution averaged across the P200 window (150-300ms) (p = 0.83). There was a significant main effect of condition in ERP distribution averaged across the P200 window (p < 0.01). There was no significant interaction effect (group x condition) effect in ERP distribution averaged across the P200 window (p = 0.42). (**B**) There was no significant man effect of group in ERP distribution averaged across the P300 window (300-500ms) (p = 0.70). There was a significant effect of condition in ERP distribution averaged across the P300 window (p < 0.01). There was no significant interaction effect (group x condition) effect in ERP distribution averaged across the P300 window (p = 0.79).

### No Difference in Theta Phase Synchronisation Power or Distribution

The TCT showed periods of topographical consistency of TPS neural activity for both groups and conditions in the time window surrounding target onset until > 400ms (see Figure S2 in supplementary materials). The 2×2 (Group x Condition) RMS randomisation test showed no significant main effect of group in TPS in the period of target onset until 900ms (global count *p* = 1, see Figure 5A). There was a significant main effect of condition from 325 to 900ms, with the attended stimuli showing higher TPS values than the non-attended stimuli (global count *p* < 0.01, see Figure 5B). There was no significant interaction effect between group and condition in this period (global count *p* = 1, see Figure 5C). The 2×2 (Group x Condition) TANOVA test showed no significant main effect of group on TPS distribution from −100 to 1000 ms surrounding target presentation (global count *p* = 1, see Figure 6A). There was a significant main effect of condition from 165 to 370 ms, indicating a different distribution of neural activity between attended and non-attended tones (global count *p* < 0.01, see Figure 6B). There was no significant interaction between group and condition (global count *p* = 0.87, see Figure 6C). The 2×2 (Group x Condition) RMS test averaged over the 150 to 300 ms after the stimulus presentation showed no significant main effect of group in TPS (*p* = 0.13). There was no significant main effect of condition (*p* = 0.74). There was no significant interaction effect of distribution between group and condition (*p* = 0.33). See Figure S3A in the supplementary materials. The 2×2 (Group x Condition) TANOVA test averaged over the 150 to 300 ms after stimulus presentation showed no significant main effect of group in TPS distribution (*p* = 0.78). There was a significant main effect of distribution between conditions (*p* < 0.001). There was no significant interaction effect of distribution between group and condition (*p* = 0.38). See Figure S3B in the supplementary materials. The 2×2 (Group x Condition) RMS test averaged over the 300 to 500 ms after stimulus presentation no significant main effect of group in TPS (*p* = 0.94). There was a significant main effect of condition (*p* < 0.02). There was no significant interaction effect of distribution between group and condition (*p* = 0.59). See Figure S4A in the supplementary materials. The 2×2 (Group x Condition) TANOVA test averaged over the 300 to 500 ms after the attended target tone showed no significant main effect of group in TPS distribution (*p* = 0.85). There was a significant main effect of distribution between conditions (*p* < 0.02). There was no significant interaction effect of distribution between group and condition (*p* = 0.99). See Figure S4B in the supplementary materials. To test whether null results were influenced by meditation experience, covariate analyses of activity from ERP and TPS measures of interest against estimated meditation experience were performed (see supplementary materials). These correlations showed meditation experience related only to RMS TPS values during the earlier window (150 to 300 ms) (p = 0.03, see Figure S6, all other p’s > 0.10).

**Fig 5.**
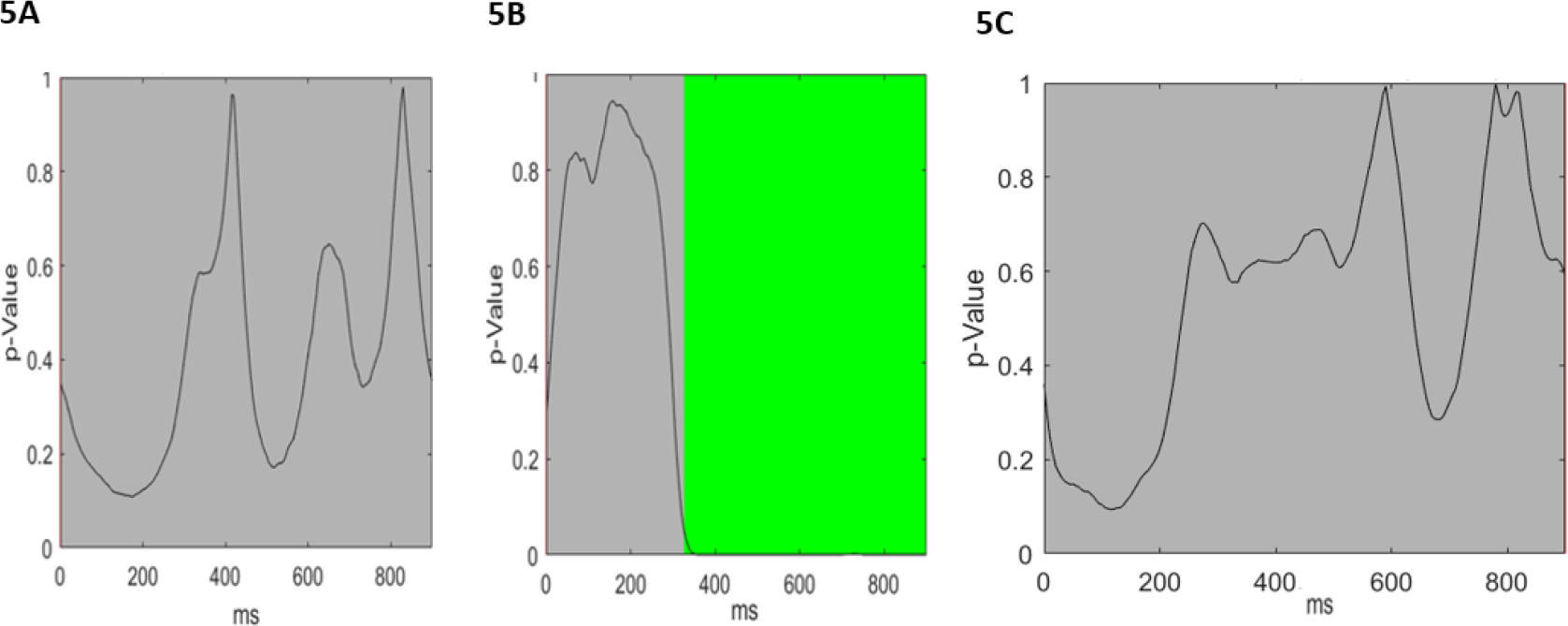
(**A**) No significant group main effect was present in TPS RMS in a period from target onset to 900ms after target onset (global count *p* = 1). The black line shows *p-*value across the epoch. (**B**) There was a significant main effect of condition in TPS across this window between attended versus not attended target trials (global count *p* < 0.01). Areas marked in green indicate the duration of a significant effect that was greater than 95% of significant effect in the randomised data, α = .05. Duration of significance within the epoch lasted from 325ms to 900ms. (**C**) There was no significant interaction effect (group x condition) (global count *p* = 1).

**Fig 6.**
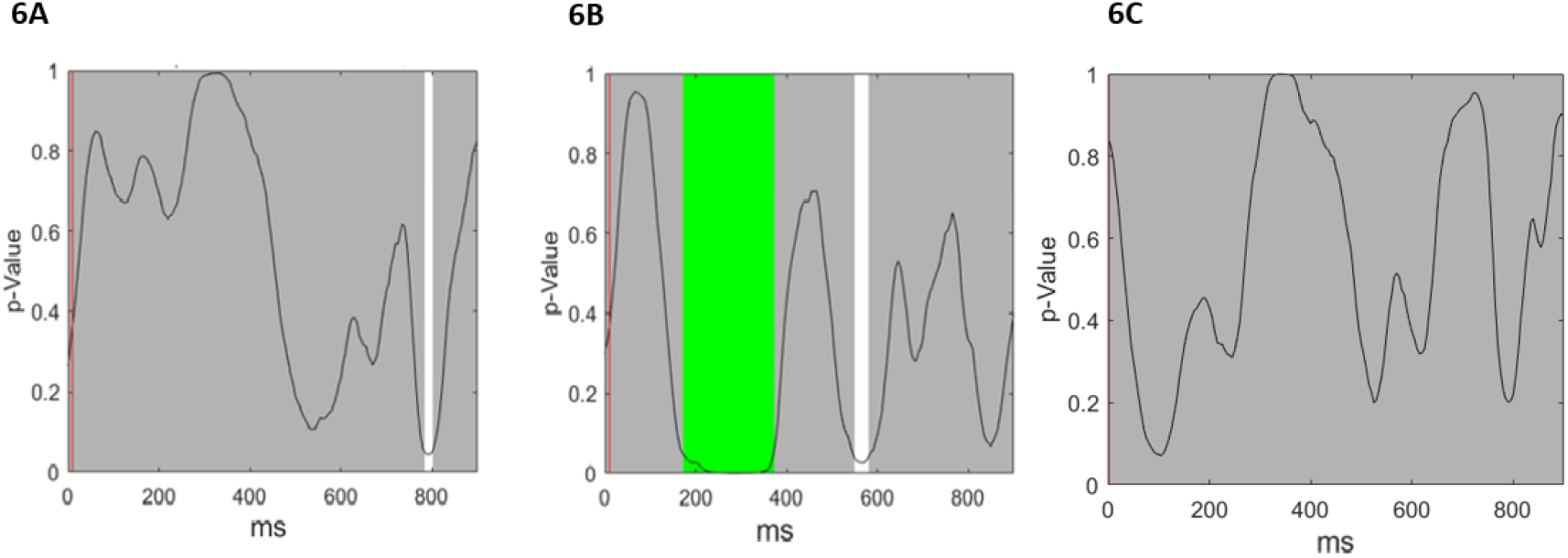
(**A**) The TANOVA revealed that there was no significant difference in distribution of TPS between groups. (**B**) The TANOVA revealed that there was a significant difference in TPS distribution between conditions lasting from 175 to 375 ms (global count *p* < 0.01). Areas marked in green indicate the duration of a significant effect that was greater 95% of significant effect in the randomised data, α = .05. (**C**) There was no significant interaction effect (group x condition) in TPS distribution (global count *p* = 0.87).

### No Correlation Between Theta Phase Synchronisation and Standard Deviation of Reaction Time

The covariate RMS randomisation test assessing the relationship between SD of RT of all participant’s and TPS during target attended trials showed no significant relationship lasting longer than duration control (both global control p = 0.10, see Figure S5 in the supplementary materials). The covariate RMS between SD of RT of participants TPS between 300 and 500ms after target attended trials showed no significant relationship when averaged across the interval (*p* = 0.52).

### Single Electrode Analyses Showed No Differences between Groups

There were no significant differences in single electrode comparisons for the P200 group comparison, nor interactions between group and attended/not attended, nor interactions between group, attended/not attended, and electrode (all p > 0.22, statistics reported in Table 3). Bayes Factor Analysis for the P200 data suggested that the null hypothesis of no differences between groups was 3.84 times more likely to be true for the alternative hypothesis, 4.15 times more likely to be true for the interaction between group and attended/not attended, and 4.38 times more likely for the interaction between group, attended/not attended and electrode. There were no significant differences in the P300 group comparison, nor interactions between group and attended/not attended, nor interactions between group, attended/not attended, and electrode (all p > 0.43, statistics reported in Table 3). Bayes Factor Analysis for the P300 data suggested that the null hypothesis of no differences between groups was 5.10 times more likely to be true for the alternative hypothesis, 2.57 times more likely to be true for the interaction between group and attended/not attended, and 7.57 times more likely for the interaction between group, attended/not attended and electrode. Similarly, there were no significant differences for TPS (150-300ms) group comparison, nor interactions between group and attended/not attended (all p > 0.09, statistics reported in Table 4). Bayes Factor Analysis for the TPS (150-300ms) comparison suggested that the null hypothesis of no differences between groups was 0.89 times more likely to be true for the alternative hypothesis, and 3.51 times more likely to be true for the interaction between group and attended/not attended. There were no significant differences for the TPS (300-500ms) group comparison, nor interactions between group and attended/not attended (all p > 0.43, statistics reported in Table 4). Bayes Factor Analysis for the TPS (300-500ms) comparison suggested that the null hypothesis of no differences between groups was 3.42 times more likely to be true for the alternative hypothesis, and 2.84 times more likely to be true for the interaction between group and attended/not attended. There was no significant correlation for TPS (300-500ms) and participants SD of RT (*p* = 0.23). Bayes Factor Analysis for the TPS (300-500ms) and SD of RT correlation suggested that the null hypothesis of no correlation was 1.49 times more likely to be true for the alternative hypothesis. Averaged activity from electrodes Fz, FCz, Cz for P200 and electrodes Fz, FCz, Cz, CPz, Pz, for P300 can be viewed in Table S1 in the supplementary materials. Averaged activity from Fz for TPS between 150-300ms and 300-500ms can be viewed in Table S2 in the supplementary materials.

**Table 3.** P200 and P300 statistics for single electrode comparisons.

|  | Null Hypothesis Test | Bayes Factor Analysis for Null Hypothesis |
| --- | --- | --- |
| **P200** |  |  |
| Between groups | $F(1,42) = 0.079, p = 0.781$, partial eta squared $< 0.001$ | 3.84 |
| Group x Attended/NotAttended | $F(1,42) = 0.523, p = 0.473$, partial eta squared $= 0.012$ | 4.15 |
| Group x Attended/NotAttended<br>x Electrode | $F(1,40) = 1.521, p = 0.224$, partial eta squared $= 0.035$ | 4.38 |
| **P300** |  |  |
| Between groups | $F(1,42) = 0.062, p = 0.805$, partial eta squared $= 0.014$ | 5.10 |
| Group x Attended/NotAttended | $F(1,42) = 0.631, p = 0.431$, partial eta squared $= 0.044$ | 2.57 |
| Group x Attended/NotAttended<br>x Electrode | $F(1,42) = 0.014, p = 0.986$, partial eta squared $= 0.003$ | 7.57 |

**Table 4.** TPS between 150-300ms and 300-500ms statistics for Fz electrode comparisons.

|  | Null Hypothesis Test | Bayes Factor Analysis for Null Hypothesis |
| --- | --- | --- |
| **TPS (150-300ms)** |  |  |
| Between groups | $F(1,42) = 3.014$, $p = 0.090$, partial eta squared = 0.067 | 0.89 |
| Group x Attended/NotAttended | $F(1,42) = 0.006$, $p = 0.941$, partial eta squared < 0.01 | 3.51 |
| **TPS (300 - 500ms)** |  |  |
| Between groups | $F(1,42) = 0.427$, $p = 0.517$, partial eta squared = 0.010 | 3.42 |
| Group x Attended/NotAttended | $F(1,42) = 0.615$, $p = 0.437$, partial eta squared = 0.014 | 2.84 |

### Power Analysis Suggested Sufficient Statistical Power

The t-test design power analysis of TPS based on the study by Lutz et al., (2009) indicated a power of 0.87, the F-test design power analysis of P300 based on Atchley et al. (2016) indicated a power of 0.98, the t-test design power analysis of P200 based on Lutz et al. (2009) indicated a power of 0.68, and the correlational design power analysis indicated power of 0.89. This suggests that the present study had the power to detect effect sizes if they were comparable to the comparisons of TPS performed by Lutz et al. (2009) and the P300 comparisons performed by Atchley et al. (2016). However, the power analysis based on the P200 from Lutz et al. (2009) suggested the study was not sufficiently powered to detect group differences if they existed. This was a surprise considering the current sample size (n = 44) was larger than Lutz et al. (2009) (n = 31). Of importance, the power analyses are based on a non-direct study design comparison with the current study, and effect sizes were obtained from traditional parametric statistics which might differ for randomisation statistics in a way that has not been clarified by previous research.

## Discussion

The current study aimed to explore whether previously identified differences in the P200, P300, and TPS are indicative of stable trait changes in a community sample of experienced meditators (in contrast to those who have recently undertaken intensive retreats and/or who are practising meditation during the task). To achieve this, an advanced electroencephalography (EEG) analysis technique compared neural response strength and distribution between a sample of healthy controls without meditation experience to experienced mindfulness meditators who were deliberately asked not to meditate during or before a dichotic listening oddball task (which was assumed to measure sustained attention).

The hypotheses that experienced meditators would display increased P200, P300, and TPS following attended target tones (compared to non-meditators) were not supported; there was no difference in neural response strength between groups. The hypotheses that the distribution of the P200, P300, and TPS would be shifted over frontal regions was also not supported. It was additionally hypothesised that there would be a negative correlation between TPS and reduced response variability; there was no significant correlation. Lastly, it was hypothesised that meditators would show higher behavioural performance than non-meditators; there was no difference in the percentage of correct target responses, d’, mean RT, or variability of RT. Power analysis revealed that the current study was sufficiently powered to detect any true differences between the groups if they existed, except for the P200 comparison (B = 0.68). However, the current sample size (n = 44) exceeded that of Lutz et al. (2009) (n = 31) who obtained significant differences in P200. Furthermore, the Bayes Factor analyses of single electrode comparisons, which replicated the comparisons from Lutz et al. (2009), suggested that the null hypothesis of no differences between groups was more likely to be true than the alternative hypothesis (except for TPS in the 150 to 300 ms time window). Bayes Factors ranged from 3.84 to 7.97 for ERP comparisons suggesting substantial evidence for the null hypothesis, and from 2.84 to 3.91 for TPS comparisons, suggesting weak to substantial evidence for the null hypothesis (Jeffereys 1961), except for the group main effect for TPS in the 150 to 300 ms window, which provided a Bayes factor of 0.89 (no evidence either way). The evidence demonstrated by these results suggests that increased P200, P300, and TPS are likely not trait changes in everyday meditators. There are several possible explanations for the null results.

### Characteristics of the Meditators

The current study’s inclusion criteria are a possible explanation of the null results. As the study aimed to observe neurophysiological changes in a community sample of meditators, the mindfulness meditation criteria were broadly defined as sitting while focusing and sustaining attention on an aspect of the present experience, such as the breath or bodily sensations. This broad inclusion criterion covers the definition of both FA and OM meditation techniques, which have been associated with unique neurophysiological changes (Cahn & Polich, 2006). The heterogeneity introduced by including either FA or OM practices might have reduced the signal to noise ratio of differences in the meditation group. Nonetheless, there were some within-group commonalities. For example, all meditators self-reported higher levels of trait mindfulness on the FFMQ than the non-meditators, suggesting that they subjectively experienced more mindful attention (though see Grossman and Van Dam (2011)). As neural activity is commonly accepted to be related to subjective experience, it would be expected that these differences would reflect changes in neural activity between groups (Tang et al., 2015). The commonalities among the meditation group were also reflected in the results of the topographical consistency test, which displayed consistent neural activity within the meditation group, suggesting common neural activity within meditators despite possible diversity of practices.

Meditation experience is also relevant to the present findings. The current sample had an approximate average meditation lifetime experience with meditation of 3,149 hours (estimated from the total years of meditation experience reported multiplied by average reported minutes meditating per week x 52 /60). While this estimate is likely to be inaccurate for a number of participants due to variability in practice across the lifetime (see Hasenkamp and Barsalou 2012), this estimate is similar to Lutz and colleagues (2009) sample who had an average of 2,967 hours. However, studies of meditators commonly include substantially more experienced participants; Delgado-Pastor et al. (2013) - approximately 5,850 hours and Cahn et al. (2013) – approximate average of 13,520 hours. The variability in meditation experience between the present study and previous work may be a potential reason for the difference in findings. Research shows that levels of meditation experience affect neural attentional network activation and the functional connectivity of distributed brain networks (Brefczynski-Lewis, Lutz, Schaefer, Levinson, & Davidson, 2007; Hasenkamp & Barsalou, 2012; Tang et al., 2015; Tomasino, Fregona, Skrap, & Fabbro, 2012). This suggestion is supported for the TPS measure at least by the correlation between TPS and estimated meditation experience. Among other possibilities, this may suggest that the minimum amount of practice for trait changes in neural patterns related to attention is closer to 5,000-10,000 hours than 3,000 hours. However, a more likely explanation is that state changes related to meditation practice during assessment; most previous research required their sample to meditate either immediately before or during their listening task.

### P200, P300, and Theta Phase Synchronisation are Not Trait Changes

In previous dichotic oddball studies, altered P200, P300, and TPS in meditators were found when meditators had been instructed to meditate immediately before the task (Cahn et al., 2013; Lutz et al., 2009; Sarang & Telles, 2006), or during the task (Atchley et al., 2016; Delgado-Pastor et al., 2013). The current study deliberately asked meditators to refrain from meditating prior to or during the target task to determine whether increased P200, P300, and TPS are neural mechanisms which reflect meditation-related trait changes resulting from long-term mindfulness meditation practice. The current study found that both groups displayed significant increases in P200, P300, and TPS when attending target tones compared to not attending target tones. This is consistent with previous research which has demonstrated that these measures relate to attentional stability (Arns et al., 2014; Lee, Lee, Kim, & Jung, 2014; Polich, 2007). However, contrary to expectations, there was no difference between meditators and non-meditators in the strength of the P200, P300, or TPS. This suggests that alterations in these mechanisms found by previous research are more likely to reflect state-related changes brought about when an experienced meditator deliberately focuses their attention rather than a generalisable trait change of mindfulness meditation. This is consistent with previous research which has reported increased theta activity, P200 and P300 amplitudes to be affected by meditation (Aftanas & Golocheikine, 2001; Cahn et al., 2013; Delgado-Pastor et al., 2013); however, research lacks consensus about what is considered a long-term trait of meditation-related neural activity (Cahn & Polich, 2006). The interpretation that increased P200, P300 amplitude, and TPS are indicative of state changes rather than trait changes may explain the present finding that meditators and non-meditators did not differ in behavioural performance on the target task. Increased TPS is proposed to aid attentional engagement and the incorporation of sensory stimuli into conscious awareness (Cahn et al., 2013; Hanslmayr et al., 2008; Kahana, Seelig, & Madsen, 2001) and increased P200 and P300 amplitudes are considered to reflect the distribution of attentional resources to incoming stimuli which facilitate information processing (Polich, 2007; Slagter et al., 2007). Thus, the groups may not have differed behaviourally because alterations in these mechanisms are related to the state of meditation, and without entering this state, experienced meditators did not generate neural activity related to higher performance than non-meditators in the dichotic listening oddball task. However, we find this explanation unlikely considering that other studies have shown experienced meditators (with comparable proficiency in meditation to the current sample) to outperform non-meditators on other attentional measures, without any meditation state changes (Bailey et al., 2018; Biedermann et al., 2016; Jha, Krompinger, & Baime, 2007). Other explanations such as the characteristics of the dichotic listening oddball are considered below.

### Characteristics of the Dichotic Listening Oddball Task

Against expectations, meditators did not perform better in the dichotic listening oddball task than controls. This is consistent with some other studies that did not include a meditation preceding or during task performance and reported no difference in behavioural performance in other attention tasks between experienced meditators and non-meditators (Josefsson & Broberg, 2011; Schmertz, Anderson, & Robins, 2009). It is possible that mindfulness meditation practice may increase attention to internal targets, such as emotions and cognitions, which may not correspond to enhanced processing of external targets. This interpretation does, however, conflict with studies assessing different measures of attention, which have shown unique neural activity and better task accuracy in experienced meditators whilst not meditating both during or prior to the task when compared to non-meditators (Bailey et al., 2018; Biedermann et al., 2016; Cardeña et al., 2015; Jha et al., 2007). One reason for this discrepancy may be the type of attention captured by the dichotic listening oddball task compared to other attentional tasks. For example, Jha et al. (2007) used the Attentional Network Test, which is a broad measure of attention including three different attentional subsystems: alerting, orienting, and conflict monitoring, whereas the dichotic listening oddball task mainly measures one aspect of attention, specifically sustained attention.

Another possibility relates to the specific dichotic listening oddball task used in the current study. While it was a direct replication of the paradigm used by Lutz et al. (2009), it was more complex, (i.e., 4-stimulus paradigm) than those of the other studies, which used simple 2-stimulus or passive paradigms (Atchley et al., 2016; Cahn et al., 2013; Delgado-Pastor et al., 2013; Jo et al., 2016; Sarang & Telles, 2006). Studies have found that simple or passive target paradigms do not always yield the same ERP and phase synchronisation measures as those produced by 3 or 4 stimulus paradigms (Choi, Cha, Choi, Jung, & Kim, 2015; Mertens & Polich, 1997). One proposed reason for this difference is that simpler paradigms only measure cognitive processes such as sensory perception and attention, whereas more complex paradigms measure these processes in addition to working memory and decision making (Choi et al., 2015); the added complexity may add noise to the neural processes being measured, reducing the signal difference between groups. Thus, the paradigm used in the present study may have compared groups behaviourally and neurophysiologically based on working memory and decision making, which may only be indirectly affected by meditation practice, in addition to sustained attention. However, there were also participants who reached perfect and close to perfect performance on the task, suggesting that a ceiling effect may have influenced behavioural results. More likely factors reducing behavioural performance may be a lower ability to discriminate the high and low tones or properly understand task instructions, which may have introduced more variability than any effect of meditation.

### Limitations

Meditators in the study were recruited based on self-reported practice rather than adherence to a specific practice for a controlled course of time. This lack of standardisation restricts the ability to draw conclusions regarding a specific meditation practice, such as OM or FA only. Stricter standardisation may be achieved through a longitudinal approach, for example testing participants before and after a mindfulness retreat. Nonetheless, while this study cannot draw conclusions about a specific meditation practice, a benefit of the cross-sectional design which included meditators who practise a variety of mindfulness techniques is that it relates to everyday mindfulness meditators in the community, who are known to change techniques throughout their life (Lea, Cadman, & Philo, 2015).

Furthermore, there were some factors which were not controlled for that may have influenced the results. One factor was that the participants’ musical and pitch discrimination abilities. As the present study used an auditory version of the task, which involved discriminating between similar sounding tones, differences in such abilities may have affected the results (although all participants included in the analysis confirmed that they could discriminate the two tones during the practice prior to performing the task). Previous studies have found that musicians are both more accurate and consistent in their responses to the auditory target task compared to non-musicians (Aschersleben, 2002), and display unique ERPs (Jongsma et al., 2005). By asking participants about their musical proficiency, participants with exceptionally high or low levels of proficiency could be excluded. Furthermore, recency of meditation practice prior to the study was not controlled for. As the state of meditation affects neurophysiology (Cahn & Polich, 2006), knowledge of when meditators had last meditated may have allowed for stronger conclusions regarding whether the studied attention-related neural activity was more reflective of state or trait neural activity.

### Implications

The present study contributes to the growing body of research that attempts to understand the neurophysiological mechanisms that underlie the effectiveness of MBIs and mindfulness meditation. The results of this study suggest that increased P200, P300, and TPS may not be long-term trait changes readily observed in a community sample of mindfulness meditators. Considering that the meditators reporting practicing mindfulness meditation for at least 2 years while the non-meditators did not, the groups likely differed qualitatively, however, this was not reflected neurophysiologically or in behavioural performance. Because subjective experience is thought to be underscored by neurophysiological processes, there are likely neurophysiological differences between these two groups which recording EEG during other attentional cognitive tasks might be able to detect. The results of this study imply that discovering trait changes in experienced meditators may be better served through focusing on different neurophysiological markers and that alterations in P200, P300, and TPS in the dichotic listening oddball task are unlikely to be generalisable trait changes in a community sample of meditators. Recommended future research can be found in the supplementary materials.

### The Main Point of the Current Research

Research attempting to understand the neural mechanisms of improved attention in mindfulness meditators is complex (Tang et al., 2015). It must account for many variables, including which attentional processes to measure and with what task, the characteristics of the meditators as well as how, and in what context, to measure their neural activity (Hölzel et al., 2011). Nonetheless, each study that evaluates a different aspect makes a valuable contribution to the developing understanding of the neural workings of mindfulness meditation (Tang et al., 2015). The present study suggests that altered P200, P300, and TPS are not trait changes resulting from mindfulness meditation practice over time. Nonetheless, discovering which trait changes result from mindfulness meditation practice is warranted as MBIs could be targeted to enhanced specific mechanisms involved in improving attention. This study shows which neurophysiological activities are not long-term markers of mindfulness meditation practice and offers direction for future researchers to pursue different neurophysiological markers with different methods of recording brain activity.

## Author Contributions

JRP performed the data collection, data analysis and wrote the paper. OB performed data collection and assisted with editing the final manuscript. HG performed data collection. BF, ME, ATH, NVD, GH, and PBF had input into study design, supported data collection or analysis, and had intellectual input and editing input into the final manuscript. NWB designed and oversaw the study, provided technical expertise and training in data analysis as well as with writing the paper.

## Compliance with Ethical Standards

All procedures performed in the study involving human participants were in accordance with the ethical standards of both The Alfred Hospital and Monash University ethical research committee and with the 1964 Helsinki declaration and its later amendments. Informed consent was obtained from all individual participants included in the study.

## Conflict of Interest and Funding

PBF has received equipment for research from MagVenture A/S, Medtronic Ltd., Cervel Neurotech and Brainsway Ltd. and funding for research from Neuronetics and Cervel Neurotech. PBF is on the scientific advisory board for Bionomics Ltd. All other authors have no conflicts to report. The study was funded by an Alfred Research Trust Small Grant Scheme (T11801). PBF is supported by a National Health and Medical Research Council of Australia Practitioner Fellowship (6069070).

## Supplementary Materials

### Electrophysiological Pre-Processing and TPS computation

The automatic procedure first rejected individual channels if more than 3% of epochs contained voltage shifts of more than +/− 250µV, a kurtosis value of >5, or power values outside of the −100 to 30 range in the 25-45 Hz band. The next step eliminated individual epochs that contained voltage shifts of more than +/− 250µV, a kurtosis value of >3, or power values outside of the −100 to 30 range in the 25-45 Hz band. Following the automatic artefact rejection, a visual inspection of the epochs ensured no artefacts remained. Independent component analysis (ICA) was performed to remove artefactual components (Chaumon, Bishop, & Busch, 2015). Adaptive Mixture ICA (AMICA) (Palmer, Makeig, Kreutz-Delgado, & Rao, 2008) was used to manually select and remove eye movements and remaining muscle activity artefacts.

After artifactual ICA components were rejected, raw data were re-filtered from 0.1-80 Hz, all previous channel and epoch rejections were applied, and rejected ICA components were applied to this 0.1-80 Hz filtered data to avoid rejecting low frequency brain activity around 1 Hz (prior to ICA rejection, data below 1 Hz was filtered out as it adversely impacts the ICA process). Rejected electrodes were re-constructed using spherical interpolation (Perrin, Pernier, Bertrand, & Echallier, 1989). Data were then visually inspected again to ensure the artefact rejection process was successful. Recordings were re-referenced offline to an averaged reference and baseline corrected to the −100 ms to 0 ms pre-stimulus period.

PLF values were calculated by first submitting EEG signal from each accepted epoch to a Morlet Wavelet Transform with a Fourier output (3.5 oscillation cycles with steps of 1 Hz and 5 ms resolution) in the theta frequency range (4-8 Hz). We then divided the Fourier spectrum by the amplitude, summed the angles, then took the absolute value of this sum and divided it by the number of trials in the specific condition and participant to obtain TPS (Delorme & Makeig, 2004).

### Assumption Tests for Participant’s Behavioural Data

Data cleaning prior to the independent samples t-test revealed no outliers in any group or measure with a z-score ± 3.29 (Tabachnick, 2013). The Shapiro-Wilk test confirmed the percentage of correct responses, SD of RT, mean RT and d’ in each group had a normal distribution. Levene’s statistic confirmed homogeneity of variance for SD of RT, mean RT, d’ and percentage of correct target responses.

### TCT Tests

**Fig S1.**
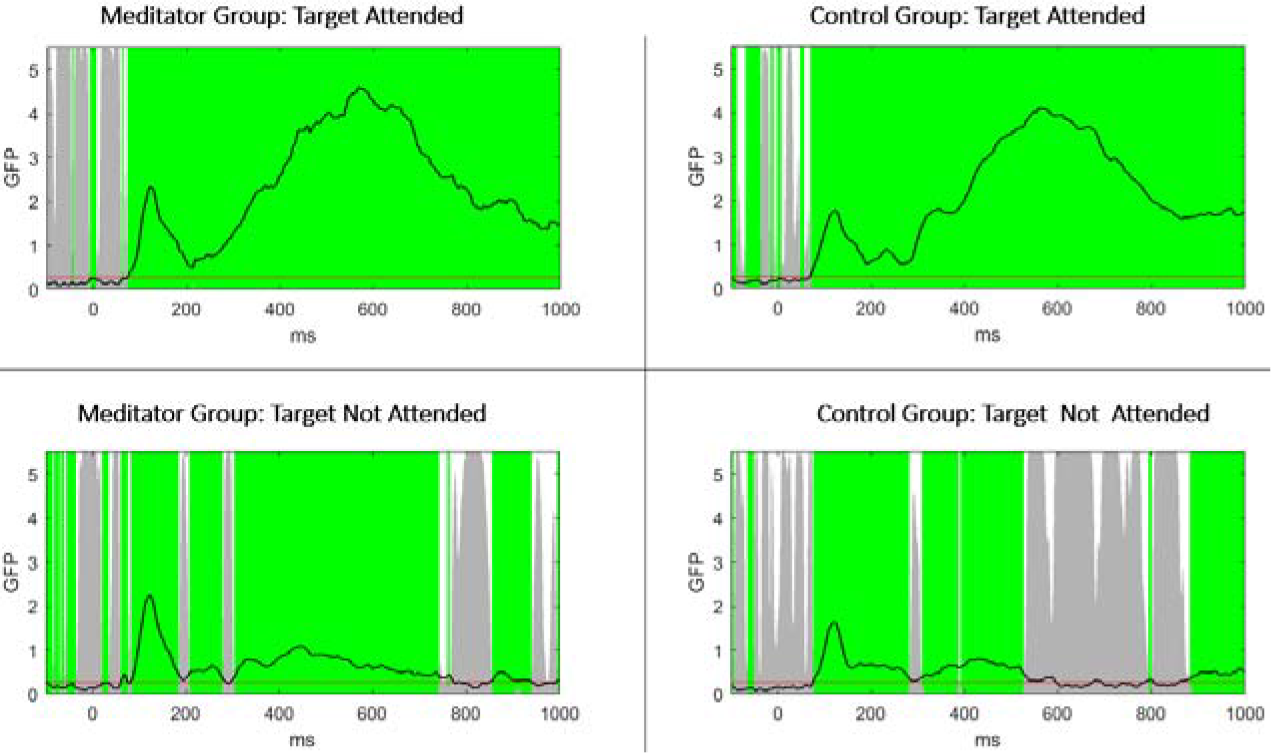
The results of the ERP TCT for each group for attended and not attended target trials. Both groups and conditions showed topographical consistency exceeding duration controls for multiple comparisons (marked in green) during the P200 window and the P300 window (300-500ms) when target was attended to, but less consistency when the target was not attended to. The horizontal red line represents a *p*-value of 0.05. Vertical grey bars represent *p* values in the TCT at each time point and the black line indicates GFP.

**Fig S2.**
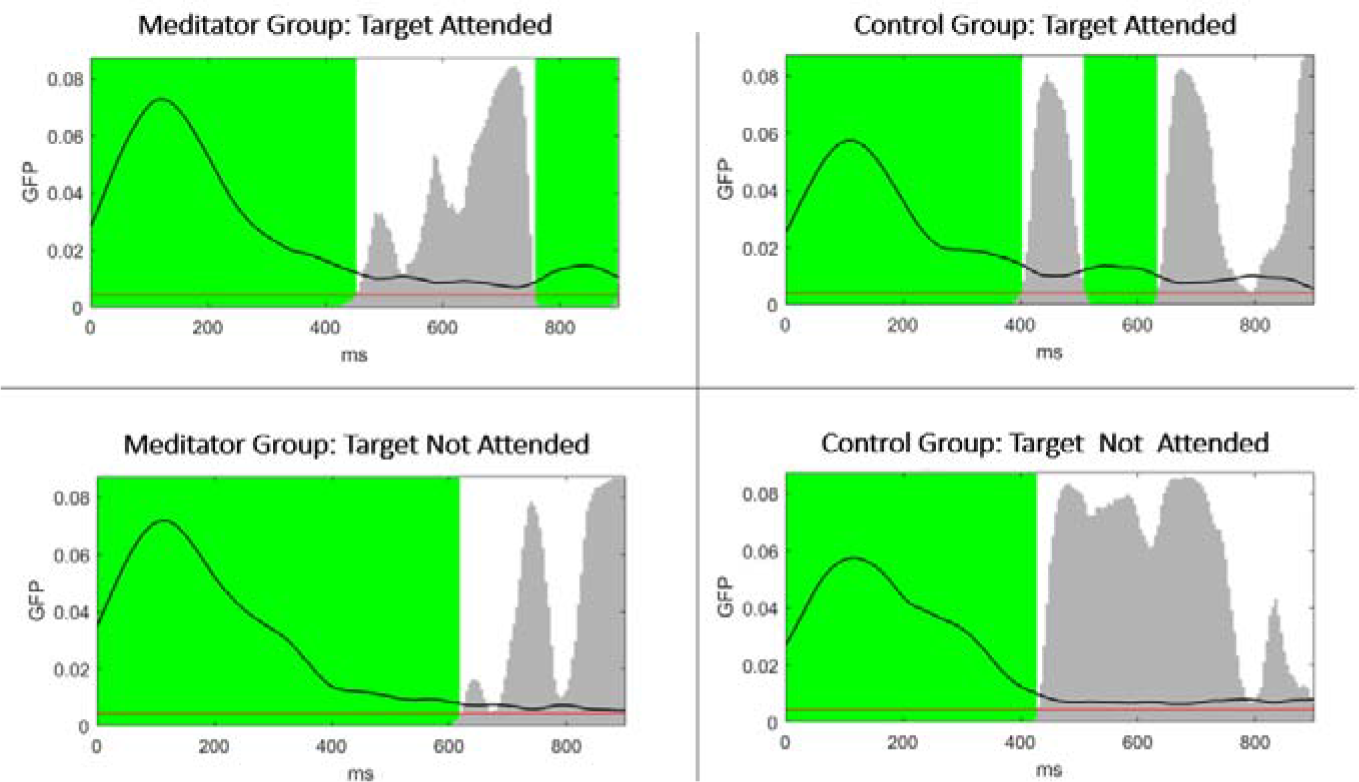
The results of the TPS TCT for each group showed periods of topographical consistency (marked in green) during the period immediately after the target stimulus was presented in both attended and not attended conditions from onset to 900ms afterwards. The red line represents a *p-*value of 0.05. Vertical grey bars represent *p* values in the TCT at each time point and the black line indicates GFP.

**Fig S3.**
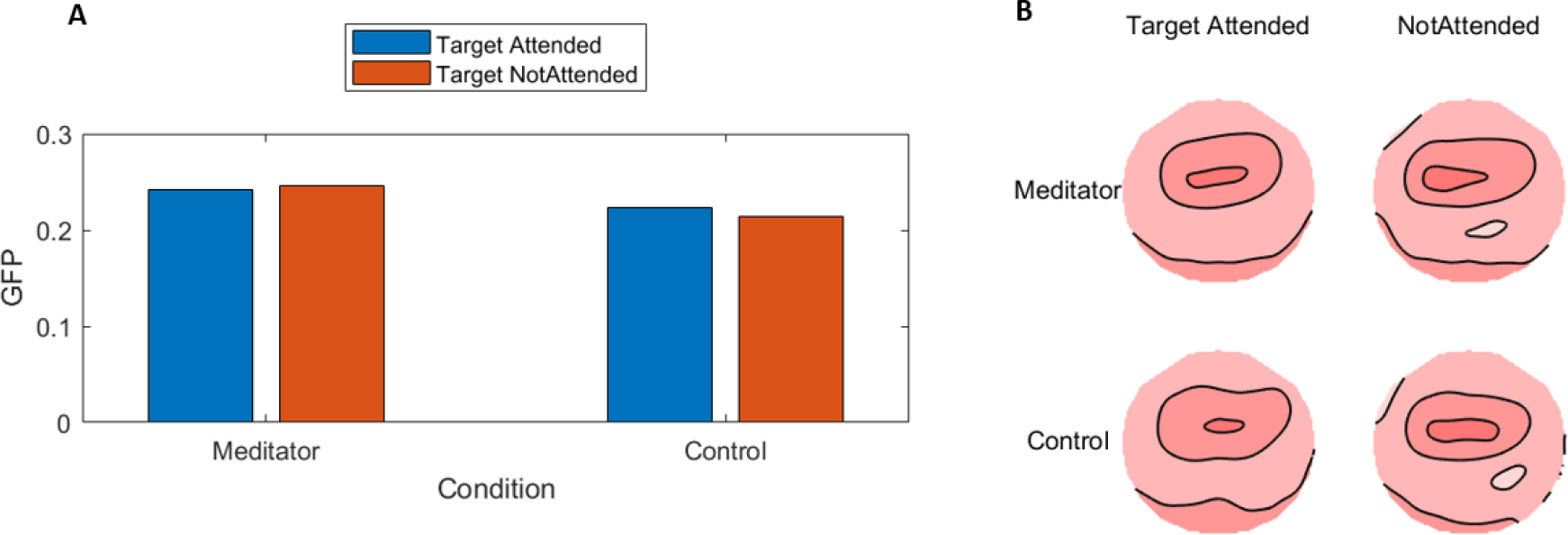
(**A**) TPS RMS test averaged over the 150 to 300 ms after stimulus presentation showed no significant main effect of group in TPS (*p* = 0.94). There was a significant main effect of condition (*p* < 0.02). There was no significant interaction effect of distribution between group and condition (*p* = 0.59). (**B**) The TANOVA test averaged over the 150 to 300 ms after the attended target tone showed no significant main effect of group in TPS distribution (*p* = 0.85). There was a significant main effect of distribution between conditions (*p* < 0.02). There was no significant interaction effect of distribution between group and condition (*p* = 0.99).

**Fig S4.**
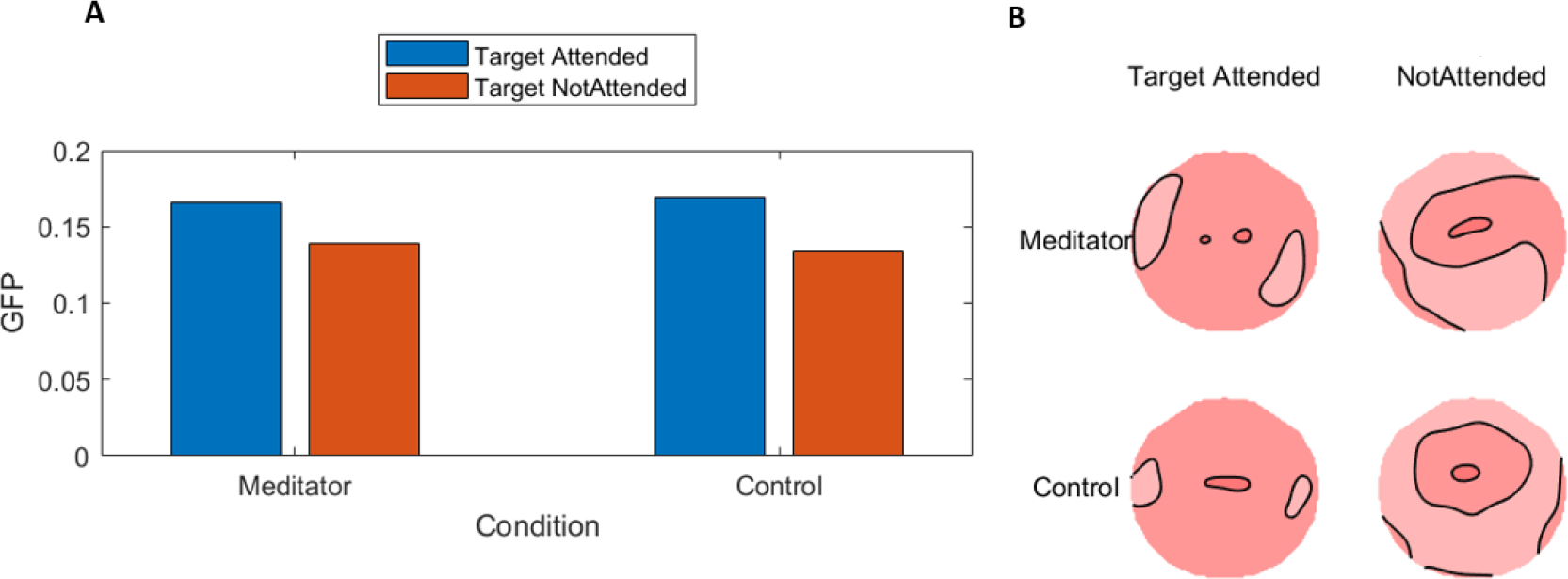
(**A**) The TPS RMS test averaged over the 300 to 500 ms after stimulus presentation showed no significant main effect of group in TPS (*p* = 0.94). There was a significant main effect of condition (*p* < 0.02). There was no significant interaction effect of distribution between group and condition (*p* = 0.59). (**B**) The TANOVA test averaged over the 300 to 500 ms after the attended target tone showed no significant main effect of group in TPS distribution (*p* = 0.85). There was a significant main effect of distribution between conditions (*p* < 0.02). There was no significant interaction effect of distribution between group and condition (*p* = 0.99).

### TPS and SD of RT

**Fig S5.**
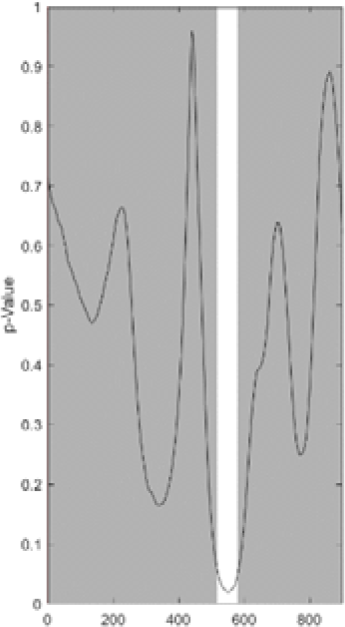
There was no significant relationship lasting longer than duration controls between RMS of TPS and the SD of RT across all participant’s during target attended trials from 0 to 900ms after the target was attended.

### Assessing the Correlation Between TPS, P200, P300 and Meditation Experience

Covariance GFP tests were conducted using estimated total meditation experience of the meditators as a covariate to measure the relationship between meditation experience and P200, P300 and a covariance RMS test assessed this relationship for TPS (150-300ms and 300-500ms). The covariate RMS randomisation test assessing the relationship between estimated total meditation experience and TPS between 150 and 300ms after target attended trials showed a significant relationship (averaged across interval p = 0.03) (See Figure S6).

**Fig S6.**
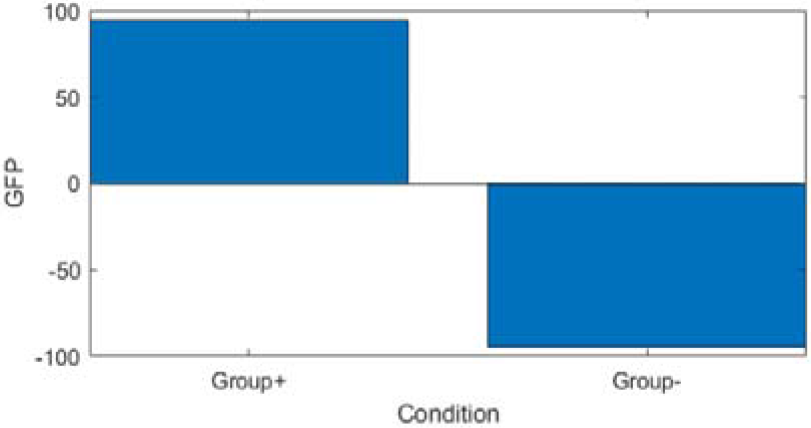
There was a relationship between the RMS of TPS (between 150 and 300ms) and estimated total meditation experience (p < 0.05 averaged across interval).

The covariate GFP randomisation test assessing the relationship between estimated total meditation experience and TPS between 300 and 500ms after target attended trials showed no significant relationship (averaged across interval p = 0.21). The covariate GFP randomisation test assessing the relationship between estimated total meditation experience and P200 (between 150 and 300ms) after target attended trials showed no significant relationship lasting longer than duration control (averaged across interval p = 0.27). The covariate GFP randomisation test assessing the relationship between estimated total meditation experience and P300 (between 300 and 500ms) after target attended trials showed no significant relationship lasting longer than duration control (averaged across interval p = 0.75).

**Table S1.**
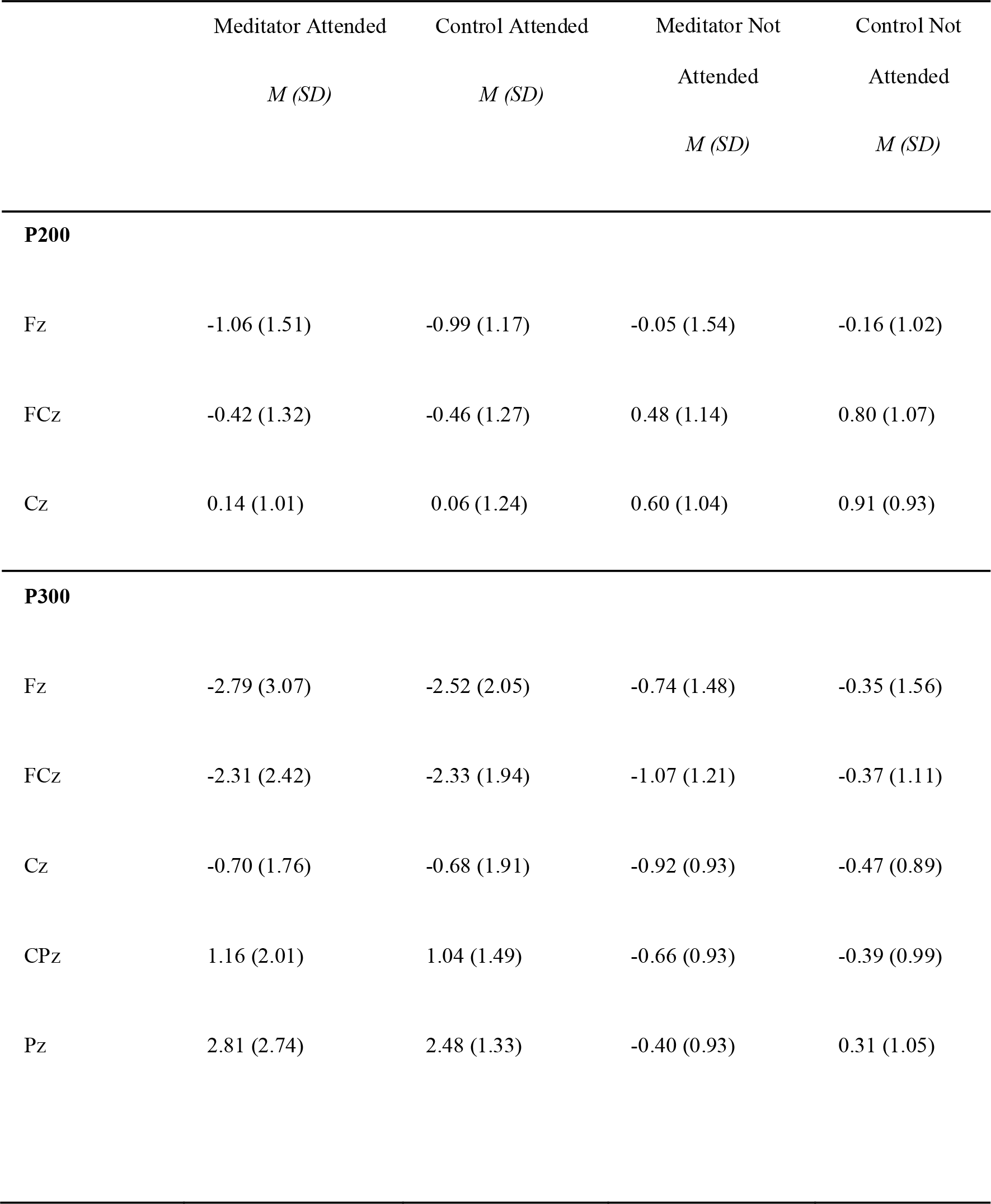
Averaged ERP activity during P200 (150-300 ms) and P300 (300-500 ms) from single electrodes.

|  | Meditator Attended | Control Attended | Meditator Not Attended | Control Not Attended |
| --- | --- | --- | --- | --- |
|  | *M (SD)* | *M (SD)* | *M (SD)* | *M (SD)* |
| **P200** |  |  |  |  |
| Fz | -1.06 (1.51) | -0.99 (1.17) | -0.05 (1.54) | -0.16 (1.02) |
| FCz | -0.42 (1.32) | -0.46 (1.27) | 0.48 (1.14) | 0.80 (1.07) |
| Cz | 0.14 (1.01) | 0.06 (1.24) | 0.60 (1.04) | 0.91 (0.93) |
| **P300** |  |  |  |  |
| Fz | -2.79 (3.07) | -2.52 (2.05) | -0.74 (1.48) | -0.35 (1.56) |
| FCz | -2.31 (2.42) | -2.33 (1.94) | -1.07 (1.21) | -0.37 (1.11) |
| Cz | -0.70 (1.76) | -0.68 (1.91) | -0.92 (0.93) | -0.47 (0.89) |
| CPz | 1.16 (2.01) | 1.04 (1.49) | -0.66 (0.93) | -0.39 (0.99) |
| Pz | 2.81 (2.74) | 2.48 (1.33) | -0.40 (0.93) | 0.31 (1.05) |

**Table S2.** Averaged TPS activity during 150-300 ms and 300-500 ms from Fz electrode.

|  | Meditator<br>Attended<br>*M (SD)* | Control Attended<br>*M (SD)* | Meditator Not<br>Attended<br>*M (SD)* | Control Not<br>Attended<br>*M (SD)* |
| --- | --- | --- | --- | --- |
| **TPS (150-300ms)** |  |  |  |  |
| Fz | 0.26 (0.10) | 0.22 (0.05) | 0.25 (0.10) | 0.21 (0.07) |
| **TPS (300-500ms)** |  |  |  |  |
| Fz | 0.17 (0.07) | 0.16 (0.05) | 0.14 (0.04) | 0.14 (0.04) |

### Future Research

Future research would benefit from pursuing several different avenues. Firstly, to further understand the context in which altered P200, P300, and TPS arise, and how specific types of meditation affect these activities. Studies could be conducted in two distinct scenarios, one in which meditators had meditated immediately prior to the task, and another when the same meditators had not. Furthermore, such studies could compare OM and FA meditators to each other, in addition to non-meditators. This may illuminate more definitively whether these neural activities are related to state or trait changes and could clarify the relationship between the state of meditation and performance during the dichotic listening task. We would recommend controlling for musical proficiency, as this is known to affect performance and neural activity during the target task (Aschersleben, 2002; Jongsma et al., 2005). Secondly, future studies using the dichotic listening oddball task should consider that the complexity of the paradigm may extend the neural activity that is measured and observed beyond simply neural processes related to attentional stability (Choi et al., 2015). This is important when comparing experienced meditators and non-meditators, as complex paradigms may not reveal the effect of long-term meditation has on the neurophysiology of attention, as they might elicit other processes (less related to meditation) as well as attention, reducing the sensitivity to potential attention-related changes that result from meditation. Thirdly, to further understand how meditation experience affects P200, P300, and TPS future studies could use a cross-sectional design with a large number of participants and a broad range of meditation experience to compare a spectrum of meditators with different amounts of meditation experience. This could reveal whether these neural activities are enhanced in more experienced meditators or if they plateau at a certain level of meditation experience. Lastly, as mindfulness meditation has become common within cognitive therapies, the study of populations suffering from disorders which involve deficits in P300, such as ADHD or depression (Dimidjian & Segal, 2015; Schoenberg et al., 2013) could be tested prior to and following an intensive mindfulness meditation program. Such a study could include three diverse types of mindfulness program, one that emphasises OM meditation, another that emphasises FA meditation, and one that incorporates both techniques. This may assist in establishing causality and specificity of technique to neural activity changes, quantifying the benefits of different mindfulness meditation practices for specific disorders, and developing an understanding as to how the mechanisms of attention can be altered to enhance MBIs.

## References

Aftanas, L. I., & Golocheikine, S. A. (2001). Human anterior and frontal midline theta and lower alpha reflect emotionally positive state and internalized attention: high-resolution EEG investigation of meditation. Neuroscience Letters, 310(1), 57–60.

Allen, M., Dietz, M., Blair, K. S., van Beek, M., Rees, G., Vestergaard-Poulsen, P., & Roepstorff, A. (2012). Cognitive-Affective Neural Plasticity following Active-Controlled Mindfulness Intervention. The Journal of Neuroscience, 32(44), 15601.

Arns, M., Jongsma, M., & Kessels, R. (2014). P300 Development across the Lifespan: A Systematic Review and Meta-Analysis. PloS One, 9(2), e87347.

Aschersleben, G. (2002). Temporal Control of Movements in Sensorimotor Synchronization. Brain and Cognition, 48(1), 66–79.

Atchley, R., Klee, D., Memmott, T., Goodrich, E., Wahbeh, H., & Oken, B. (2016). Event-related potential correlates of mindfulness meditation competence. Neuroscience, 320, 83–92.

Badart, P., McDowall, J., & Prime, S. (2018). Multimodal Sustained Attention Superiority in Concentrative Meditators Compared to Nonmeditators. Mindfulness, 9(3), 824–835.

Baer, R. A., Smith, G. T., Hopkins, J., Krietemeyer, J., & Toney, L. (2006). Using Self-Report Assessment Methods to Explore Facets of Mindfulness. Assessment, 13(1), 27–45.

Baijal, S., & Srinivasan, N. (2010). Theta activity and meditative states: spectral changes during concentrative meditation. International Quarterly of Cognitive Science, 11(1), 31–38.

Bailey, N., Freedman, G., Raj, K., Sullivan, C., Rogasch, N., Chung, S. W., & Fitzgerald, P. (2018). Mindfulness meditators show altered distributions of early and late neural activity markers of attention in a response inhibition task. bioRxiv, 396259.

Bailey, N.W., Raj, K., Freedman., Fitzgibbon, B., Rogasch, N. C., Van Dam, N., & Fitzgerald, P. (2019). Mindfulness meditators do not show differences in electrophysiological measures of error processing. Mindfulness, 10(2), 1–21.

Baird, B., Smallwood, J., Lutz, A., & Schooler, J. W. (2014). The Decoupled Mind: Mind-wandering Disrupts Cortical Phase-locking to Perceptual Events. Journal of Cognitive Neuroscience, 26(11), 2596–2607.

Barron, E., Riby, L. M., Greer, J., & Smallwood, J. (2011). Absorbed in Thought: The Effect of Mind Wandering on the Processing of Relevant and Irrelevant Events. Psychological Science, 22(5), 596–601.

Beck, A. T., Steer, R. A., & Brown, G. K. (1996). Beck depression inventory-II. San Antonio, 78(2), 490–498.

Biedermann, B., de Lissa, P., Mahajan, Y., Polito, V., Badcock, N., Connors, M. H., … McArthur, G. (2016). Meditation and auditory attention: An ERP study of meditators and non-meditators. International Journal of Psychophysiology, 109(C), 63–70.

Brefczynski-Lewis, J. A., Lutz, A., Schaefer, H. S., Levinson, D. B., & Davidson, R. J. (2007). Neural correlates of attentional expertise in long-term meditation practitioners. Proceedings of the National Academy of Sciences of the United States, 104(27), 11483.

Britton, W. B., Davis, J. H., Loucks, E. B., Peterson, B., Cullen, B. H., Reuter, L., & Lindahl, J. R. (2018). Dismantling Mindfulness-Based Cognitive Therapy: Creation and validation of 8-week focused attention and open monitoring interventions within a 3-armed randomized controlled trial. Behaviour Research and Therapy, 101, 92–107.

Cahn, Delorme, A., & Polich, J. (2013). Event-related delta, theta, alpha and gamma correlates to auditory oddball processing during Vipassana meditation. Social Cognitive and Affective Neuroscience, 8(1), 100–111.

Cahn, & Polich, J. (2006). Meditation states and traits: EEG, ERP, and neuroimaging studies. Psychological Bulletin, 132(2), 180–211.

Cahn, & Polich, J. (2009). Meditation (Vipassana) and the P3a event-related brain potential. International Journal of Psychophysiology, 72(1), 51–60.

Cardeña, E., Sjöstedt, J., & Marcusson-Clavertz, D. (2015). Sustained Attention and Motivation in Zen Meditators and Non-meditators. Mindfulness, 6(5), 1082–1087.

Chambers, R., Gullone, E., & Allen, N. B. (2009). Mindful emotion regulation: An integrative review. Clinical Psychology Review, 29(6), 560–572.

Chaumon, M., Bishop, D. V. M., & Busch, N. A. (2015). A practical guide to the selection of independent components of the electroencephalogram for artifact correction. Journal of Neuroscience Methods, 250, 47–63.

Choi, J. W., Cha, K. S., Choi, J. D., Jung, K.-Y., & Kim, K. H. (2015). Difficulty-related changes in inter-regional neural synchrony are dissociated between target and non-target processing. Brain Research, 1603(C), 114–123.

Davidson, R. (2005). Meditation and Neuroplasticity: Training Your Brain. Explore: The Journal of Science and Healing, 1(5), 380–388.

Delgado-Pastor, L. C., Perakakis, P., Subramanya, P., Telles, S., & Vila, J. (2013). Mindfulness (Vipassana) meditation: Effects on P3b event-related potential and heart rate variability. International Journal of Psychophysiology, 90(2), 207–214.

Delorme, A., & Makeig, S. (2004). EEGLAB: an open source toolbox for analysis of single-trial EEG dynamics including independent component analysis. Journal of Neuroscience Methods, 134(1), 9–21.

Faul, F., Erdfelder, E., & Buchner, A. (2007). G*Power 3: A flexible statistical power analysis program for the social, behavioral, and biomedical sciences. Behavior Research Methods, 39(2), 175–191.

Godfrin, K. A., & van Heeringen, C. (2010). The effects of mindfulness-based cognitive therapy on recurrence of depressive episodes, mental health and quality of life: A randomized controlled study. Behaviour Research and Therapy, 48(8), 738–746.

Goyal, M., Singh, S., Sibinga, E. S., & et al. (2014). Meditation programs for psychological stress and well-being: A systematic review and meta-analysis. JAMA Internal Medicine, 174(3), 357–368.

Grant, J. A., Courtemanche, J., Duerden, E. G., Duncan, G. H., & Rainville, P. (2010). Cortical thickness and pain sensitivity in zen meditators. Emotion (Washington, D.C.), 10(1), 43.

Greenhouse, S. W., & Geisser, S. (1959). On methods in the analysis of profile data. Psychometrika, 24(2), 95–112.

Grossman, P., & Van Dam, N. T. (2011). Mindfulness, by any other name…: trials and tribulations of sati in western psychology and science. Contemporary Buddhism, 12(1), 219–239.

Gu, J., Strauss, C., Bond, R., & Cavanagh, K. (2015). How do mindfulness-based cognitive therapy and mindfulness-based stress reduction improve mental health and wellbeing? A systematic review and meta-analysis of mediation studies. Clinical Psychology Review, 37, 1–12.

Habermann, M., Weusmann, D., Stein, M., & Koenig, T. (2018). A Student's Guide to Randomization Statistics for Multichannel Event-Related Potentials Using Ragu. Frontiers in Neuroscience, 12.

Hanslmayr, S., Pastötter, B., Bäuml, K.-H., Gruber, S., Wimber, M., & Klimesch, W. (2008). The Electrophysiological Dynamics of Interference during the Stroop Task. Journal of Cognitive Neuroscience, 20(2), 215–225.

Hasenkamp, W., & Barsalou, L. W. (2012). Effects of meditation experience on functional connectivity of distributed brain networks. Frontiers in Human Neuroscience, 6(2012), 1–14.

Hergueta, T., Baker, R., & Dunbar, G. C. (1998). The Mini-International Neuropsychiatric Interview (MINI): the development and validation of a structured diagnostic psychiatric interview for DSM-IVand ICD-10. Journal of Clinical Psychiatry, 59(20), 2233.

Hodgins, H. S., & Adair, K. C. (2010). Attentional processes and meditation. Consciousness and Cognition, 19(4), 872–878.

Hölzel, B. K., Lazar, S. W., Gard, T., Schuman-Olivier, Z., Vago, D. R., & Ott, U. (2011). How Does Mindfulness Meditation Work? Proposing Mechanisms of Action From a Conceptual and Neural Perspective. Perspectives on Psychological Science, 6(6), 537–559.

Jeffreys, H. (1961). Theory of probability (3rd ed.). Oxford: Clarendon Press.

Jha, A. P., Krompinger, J., & Baime, M. J. (2007). Mindfulness training modifies subsystems of attention. Cognitive, Affective, & Behavioral Neuroscience, 7(2), 109–119.

Jo, H.-G., Schmidt, S., Inacker, E., Markowiak, M., & Hinterberger, T. (2016). Meditation and attention: A controlled study on long-term meditators in behavioral performance and event-related potentials of attentional control. International Journal of Psychophysiology, 99, 33–39.

Jongsma, M. L. A., Eichele, T., Quiroga, R. Q., Jenks, K. M., Desain, P., Honing, H., & Van Rijn, C. M. (2005). Expectancy effects on omission evoked potentials in musicians and non musicians. Psychophysiology, 42(2), 191–201.

Josefsson, T., & Broberg, A. (2011). Meditators and non-meditators on sustained and executive attentional performance. Mental Health, Religion & Culture, 14(3), 291–309.

Kabat-Zinn, J. (1994). Wherever you go. There you are: mindfulness meditation in everyday life. New York: Hyperion.

Kahana, M. J., Seelig, D., & Madsen, J. R. (2001). Theta returns. Current Opinion in Neurobiology, 11(6), 739–744.

Keng, S.-L., Smoski, M. J., & Robins, C. J. (2011). Effects of mindfulness on psychological health: A review of empirical studies. Clinical Psychology Review, 31(6), 1041–1056.

Klimesch, Doppelmayr, M., Schimke, H., & Ripper, B. (1997). Theta synchronization and alpha desynchronization in a memory task. Psychophysiology, 34(2), 169–176.

Klimesch, Sauseng, P., Hanslmayr, S., Gruber, W., & Freunberger, R. (2007). Event-related phase reorganization may explain evoked neural dynamics. Neuroscience and Biobehavioral Reviews, 31(7), 1003–1016.

Koenig, T., Kottlow, M., Stein, M., & Melie-García, L. (2011). Ragu: A Free Tool for the Analysis of EEG and MEG Event-Related Scalp Field Data Using Global Randomization Statistics. Computational Intelligence and Neuroscience, 2011(2011), 14.

Koenig, T., & Melie-garcía, L. (2010). A Method to Determine the Presence of Averaged Event-Related Fields Using Randomization Tests. Brain Topography, 23(3), 233–242.

Lakey, C. E., Berry, D. R., & Sellers, E. W. (2011). Manipulating attention via mindfulness induction improves p300-based braincomputer interface performance. Journal of Neural Engineering, 8(2), 025019.

Lea, J., Cadman, L., & Philo, C. (2015). Changing the habits of a lifetime? Mindfulness meditation and habitual geographies. Cultural Geographies, 22(1), 49–65.

Lee, G.-T., Lee, C., Kim, K. H., & Jung, K.-Y. (2014). Regional and inter-regional theta oscillation during episodic novelty processing. Brain and Cognition, 90, 70–75.

Light, G. A., Williams, L. E., Minow, F., Sprock, J., Rissling, A., Sharp, R., & Braff, D. L. (2010). Electroencephalography (EEG) and event-related potentials (ERPs) with human participants. Current Protocols in Neuroscience, 6(52), Unit 6.25.21.

Lindholm, E., & Koriath, J. J. (1985). Analysis of multiple event related potential components in a tone discrimination task. International journal of psychophysiology : official journal of the International Organization of Psychophysiology, 3(2), 121.

Lutz, Slagter, Dunne, & Davidson. (2008). Attention regulation and monitoring in meditation. Trends in Cognitive Sciences, 12(4), 163–169.

Lutz, Slagter, H., A, Rawlings, N., B, Francis, D., A, Greischar, L., L, & Davidson, J., D. (2009). Mental Training Enhances Attentional Stability: Neural and Behavioral Evidence. The Journal of Neuroscience, 29(42), 13418–13427.

Ma, S. H., & Teasdale, J. D. (2004). Mindfulness-based cognitive therapy for depression: replication and exploration of differential relapse prevention effects. Journal of Consulting and Clinical Psychology, 72(1), 31.

Mertens, R., & Polich, J. (1997). P300 from a single-stimulus paradigm: passive versus active tasks and stimulus modality. Electroencephalography and Clinical Neurophysiology, 104(6), 488.

Miller, J. J., Fletcher, K., & Kabat-Zinn, J. (1995). Three-year follow-up and clinical implications of a mindfulness meditation-based stress reduction intervention in the treatment of anxiety disorders. General Hospital Psychiatry, 17(3), 192–200.

Oostenveld, R., Fries, P., Maris, E., & Schoffelen, J.-M. (2011). FieldTrip: Open Source Software for Advanced Analysis of MEG, EEG, and Invasive Electrophysiological Data. Computational Intelligence and Neuroscience, 2011(2011), 9.

Palmer, J., Makeig, S., Delgado, K., & Rao, B. (2008). Newton method for the ICA mixture model. 2008 IEEE International Conference on Acoustics, Speech and Signal Processing, 1805–1808.

Perrin, F., Pernier, J., Bertrand, O., & Echallier, J. (1989). Spherical splines for scalp potential and current density mapping. Electroencephalography and clinical neurophysiology, 72(2), 184–187.

Polich, J. (2007). Updating P300: An integrative theory of P3a and P3b. Clinical Neurophysiology, 118(10), 2128–2148.

Rouder, J. N., Morey, R. D., Verhagen, J., Swagman, A. R., & Wagenmakers, E.-J. (2017). Bayesian analysis of factorial designs. Psychological Methods, 22(2), 304–321.

Sarang, S. P., & Telles, S. (2006). CHANGES IN P300 FOLLOWING TWO YOGA-BASED RELAXATION TECHNIQUES. International Journal of Neuroscience, 116(12), 1419–1430.

Schmertz, S., Anderson, P., & Robins, D. (2009). The Relation Between Self-Report Mindfulness and Performance on Tasks of Sustained Attention. Journal of Psychopathology and Behavioral Assessment, 31(1), 60–66.

Schoenberg, P. L., & Vago, D. R. (2019). Mapping meditative states and stages with electrophysiology: concepts, classifications, and methods. Current Opinion in Psychology, 28, 211–217.

Slagter, H. A., Lutz, A., Greischar, L. L., Francis, A. D., Nieuwenhuis, S., Davis, J. M., & Davidson, R. J. (2007). Mental Training Affects Distribution of Limited Brain Resources. PLoS Biology, 5(6), e138.

Slagter, H. A., Lutz, A., Greischar, L. L., Nieuwenhuis, S., & Davidson, R. J. (2009). Theta Phase Synchrony and Conscious Target Perception: Impact of Intensive Mental Training. Journal of Cognitive Neuroscience, 21(8), 1536–1549.

Steer, R. A., & Beck, A. T. (1997). Beck Anxiety Inventory.

Tang, Hölzel, B. K., & Posner, M. I. (2015). The neuroscience of mindfulness meditation. Nature Reviews. Neuroscience, 16(4), 213–225.

Thomas, J. W., & Cohen, M. (2014). A methodological review of meditation research. Frontiers in psychiatry, 5, 74–74.

Tomasino, B., & Fabbro, F. (2016). Increases in the right dorsolateral prefrontal cortex and decreases the rostral prefrontal cortex activation after-8 weeks of focused attention based mindfulness meditation. Brain and Cognition, 102, 46–54.

Tomasino, B., Fregona, S., Skrap, M., & Fabbro, F. (2012). Meditation-related activations are modulated by the practices needed to obtain it and by the expertise: an ALE meta-analysis study. Frontiers in Human Neuroscience, 6, 346.

Van Dam, N. T., van Vugt, M. K., Vago, D. R., Schmalzl, L., Saron, C. D., Olendzki, A., & Meyer, D. E. (2018). Mind the Hype: A Critical Evaluation and Prescriptive Agenda for Research on Mindfulness and Meditation. Perspectives on Psychological Science, 13(1), 36–61.

